# Low-dimensional Dynamics of Two Coupled Biological Oscillators

**DOI:** 10.1101/541128

**Authors:** Colas Droin, Eric R Paquet, Felix Naef

## Abstract

The circadian clock and the cell cycle are two biological oscillatory processes that coexist within individual cells. These two oscillators were found to interact, which can lead to their synchronization. Here, we develop a method to infer their coupling and non-linear dynamics from thousands of mouse and human single-cell microscopy traces. This coupling predicts multiple phase-locked states showing different degrees of robustness against molecular fluctuations inherent to cellular scale biological oscillators. Moreover, the phase–locked states were temperature-independent and evolutionarily conserved from mouse to human, hinting at a common underlying dynamical mechanism. Finally, we detected a signature of the coupled dynamics in a physiological context, where tissues with different proliferation states exhibited shifted circadian clock phases.

## Introduction

The circadian clock and the cell cycle are two periodic processes that cohabit in many types of living cells. In single mammalian cells, circadian clocks consist of autonomous feedback loop oscillators ticking with an average period of about 24h^1^, and controlling many downstream cellular processes^2^. In conditions of high proliferation such as those found in cultured cells or certain tissues, the cell-cycle progresses essentially continuously and can thus be abstracted as an oscillator with an average period matching the cell doubling time. Both processes fluctuate due to intra-cell molecular noise, as well as external fluctuations. While the precision of the circadian period is typically about 15% in fibroblast cells^1^, the cell cycle can be more variable depending on the conditions and cell lines^3,4^. Interestingly, previous work showed that the two cycles can mutually interact^1^, which may then lead, as theory predicts, to synchronized dynamics^5,6^ and important physiological consequences such as cell-cycle synchrony during liver regeneration^7^. In tissue-culture cells, which are amenable to systematic microscopy analysis, it was found that the phase dynamics of two oscillators shows phase-locking^5,6^, defined by a rational rotation number *p*: *q* such that *p* cycles of one oscillator are completed while the other completes *q*8.

Concerning the nature of those interactions, the influence of the circadian clock on cell-cycle progression and division timing has been shown in several systems^7,9–13^. In contrast, we showed in mouse fibroblasts that the cell cycle strongly influences the circadian oscillator^5^, which was also investigated theoretically and linked with DNA replication in bacteria^14^. In addition, human cells can switch between a state of high cell proliferation with a damped circadian oscillator, to a state of low proliferation but robust circadian rhythms, depending on molecular interactions and activities of cell cycle and clock regulators^15^.

Here, we exploit the fact that the two coupled cycles evolve on a low dimensional and compact manifold (the flat torus) to fully characterize their dynamics. In particular, starting from a generic stochastic model for the interacting phases combined with fluorescence microscopy recordings from thousands of individual cells, we obtained a data-driven reconstruction of the coupling function describing how the cell cycle influences the circadian oscillator. This coupling phase-locks the two oscillators in a temperature-independent manner, and only few of the deterministically predicted phase-locked states were stable against inherent fluctuations. Moreover, we established that the coupling between the two oscillators is conserved from mouse to human, and can override systemic synchronization signals such as temperature cycles. Finally, we showed in a physiological context how this coupling explains why mammalian tissues with different cell proliferation rates have shifted circadian phases.

## Results

### A data-driven reconstruction of the phase dynamics and coupling of two biological oscillators

To study the phase dynamics of the circadian and cell cycle oscillators, we reconstructed a stochastic dynamic model of the two coupled oscillators from single-cell time-lapse microscopy traces of a fluorescent Rev-Erb-α-YFP circadian reporter^1,5^.

#### Phase model

First, we represent the phase dynamics of the circadian oscillator (*θ* = 0 corresponds to peaks of fluorescence) and cell-cycle (*φ* = 0 is the cytokinesis) on a 0,2*π*)×[0,2*π*) torus. Since we showed previously that the influence of the clock on the cell cycle was negligible in 3T3 cells^5^, we here model only how the cell-cycle progression influences the instantaneous circadian phase velocity *ω*_*θ*_ using a general coupling function *F*(*θ*, *φ*) (Fig. 1a). To account for circadian phase fluctuations and variability in circadian period length known to be present in single cells^1,16^, we added a phase diffusion term *σ*_*θ*_*dW*^17,18^. For the cell-cycle phase, we assumed a piecewise linear and deterministic phase progression in between two successive divisions. The phase model reads:

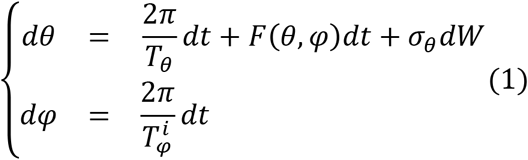

**Figure 1:**
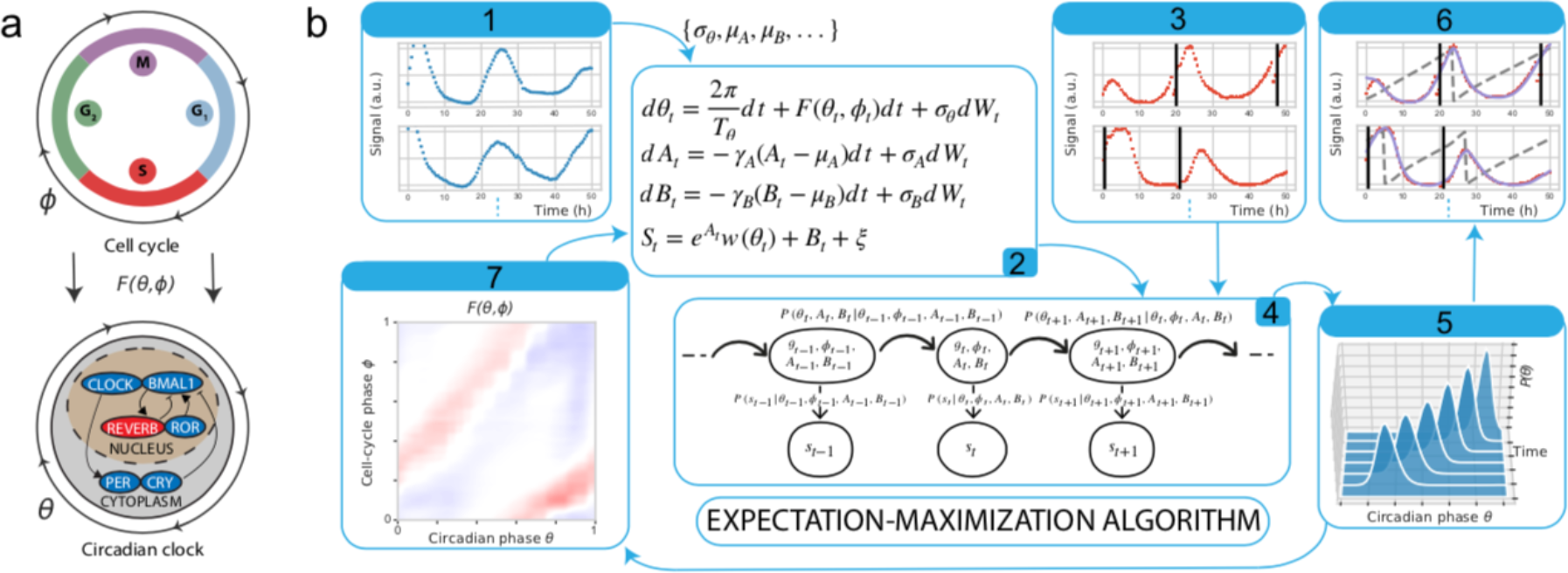
Reconstructing the phase dynamics and coupling of two biological oscillators. (**a**) In mouse fibroblasts, the cell cycle (top) can influence the circadian oscillator (bottom) according to a coupling function *F*(*θ*, *φ*), where *φ* denote the cell-cycle and *θ* the circadian oscillator phases. (**b**) Fluorescence microscopy traces (Rev-Erb-α-YFP circadian reporter) are recorded for non-dividing and dividing cells (boxes 1 and 3, respectively). These signals *S*_*t*_ are explicitly modeled (box 2) using diffusion-drift equations for the circadian phase *θ*_*t*_, amplitude *A*_*t*_ and background *B*_*t*_ fluctuations, as well with a function *w* to map the phase *θ*_*t*_ to the observation space. The coupling function *F*(*θ*, *φ*) enters the equation for *θ* only in the case of dividing cells. Most parameters are estimated from the non-dividing cells, while the coupling function *F*(*θ*, *φ*) is inferred using a maximum likelihood formulation. The optimization problem is solved by converting to a HMM (box 4) in which *θ*_*t*_, *A*_*t*_ and *B*_*t*_ are latent variables. The HMM is used on traces to compute posterior probabilities of circadian phases (box 5), while the cell-cycle phase is retrieved using linear interpolation between successive divisions (3, vertical black lines). Expected circadian phases (dashed grey line) and fits (purple line) to the data (red) are shown in box 6. An iterative EM algorithm then yields the converged coupling *F*(*θ*, *φ*)(box 7).

Here, *T*_*θ*_ represents the intrinsic circadian period, while the term 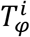 represents the *i*^*th*^ cell-cycle interval between two successive divisions. The goal will be to identify the function

*F*(*θ*, *φ*) and study the consequences on the coupled oscillator dynamics.

#### Model of the signal

We linked the circadian phase with the measured time traces through a 2*π*-periodic function *w*(*θ*). In addition, as indicated by typical data traces (Supplementary Fig. 1a), we considered amplitude (*A*_*t*_) and baseline (*B*_*t*_) fluctuations, which for simplicity we modeled as independent from *θ*_*t*_, an assumption that was supported *a posteriori* (Supplementary Information). The full model for the signal *S*_*t*_ thus reads:

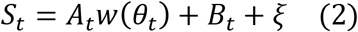

where *ξ* is a normally distributed random variable (measurement noise) and *A*_*t*_, *B*_*t*_ are Ornstein-Uhlenbeck processes varying more slowly than the phase distortion caused by *F*(*θ*, *φ*), *i.e.*, on timescales on the order of the circadian period (Supplementary Information).

#### Inference of phases & coupling function

From this stochastic model (Equations 1&2), we built a Hidden Markov Model (HMM) to calculate posterior probabilities of the oscillator phases at each measured time point, using the forward-backward algorithm^19^. To estimate *F*(*θ*, *φ*), we used a maximum-likelihood approach that combines goodness of fit with sparseness and smoothness constraints, which we implemented with an Expectation-Maximization (EM) algorithm (Methods, Supplementary Information).

The successive steps of our approach are illustrated in Figure 1b. Dividing cells indicated that, typically, the circadian phase progression shows variations in phase velocity (Supplementary Fig. 1a). To validate that these variations can identify *F*(*θ*, *φ*), we generated noisy traces *in silico* with pre-defined *F*(*θ*, *φ*) and reconstructed the coupling function, showing excellent qualitative agreement (Supplementary Fig. 1b-c). Nevertheless, the magnitude of the coupling tended to be underestimated, presumably owing to the stochasticity of the system, observation noise and regularization terms used.

### Influence of the cell cycle on the circadian phase progression

In mouse embryonic fibroblasts (NIH3T3), we showed that due to the coupling, circadian periods decrease with temperature in dividing cells, but not in quiescent cells^5^. To further understand how temperature influences the interaction between the two oscillators, we re-analyzed NIH3T3 traces (24-72h long) obtained at 34°C, 37°C, and 40°C^5^. From those, we found that both the inferred coupling functions and phase densities at the three temperatures were very similar, with almost identical 1:1 phase-locked orbits (Supplementary Fig. 2a-c). We therefore modeled the coupling as temperature-independent and re-constructed a definitive *F*(*θ*, *φ*) from traces at all temperatures (Fig. 2a, Supplementary Fig. 2d). Unlike the localized coupling assumed in our previous work^5^, this function shows a diffuse structure mainly composed of two juxtaposed diagonal stripes: one for phase acceleration (red), and one, less structured, for deceleration (blue). The slopes of these stripes are about one, however, the phase velocity varies along the stripes, which justifies using a 2D parameterization of the coupling function.

**Figure 2:**
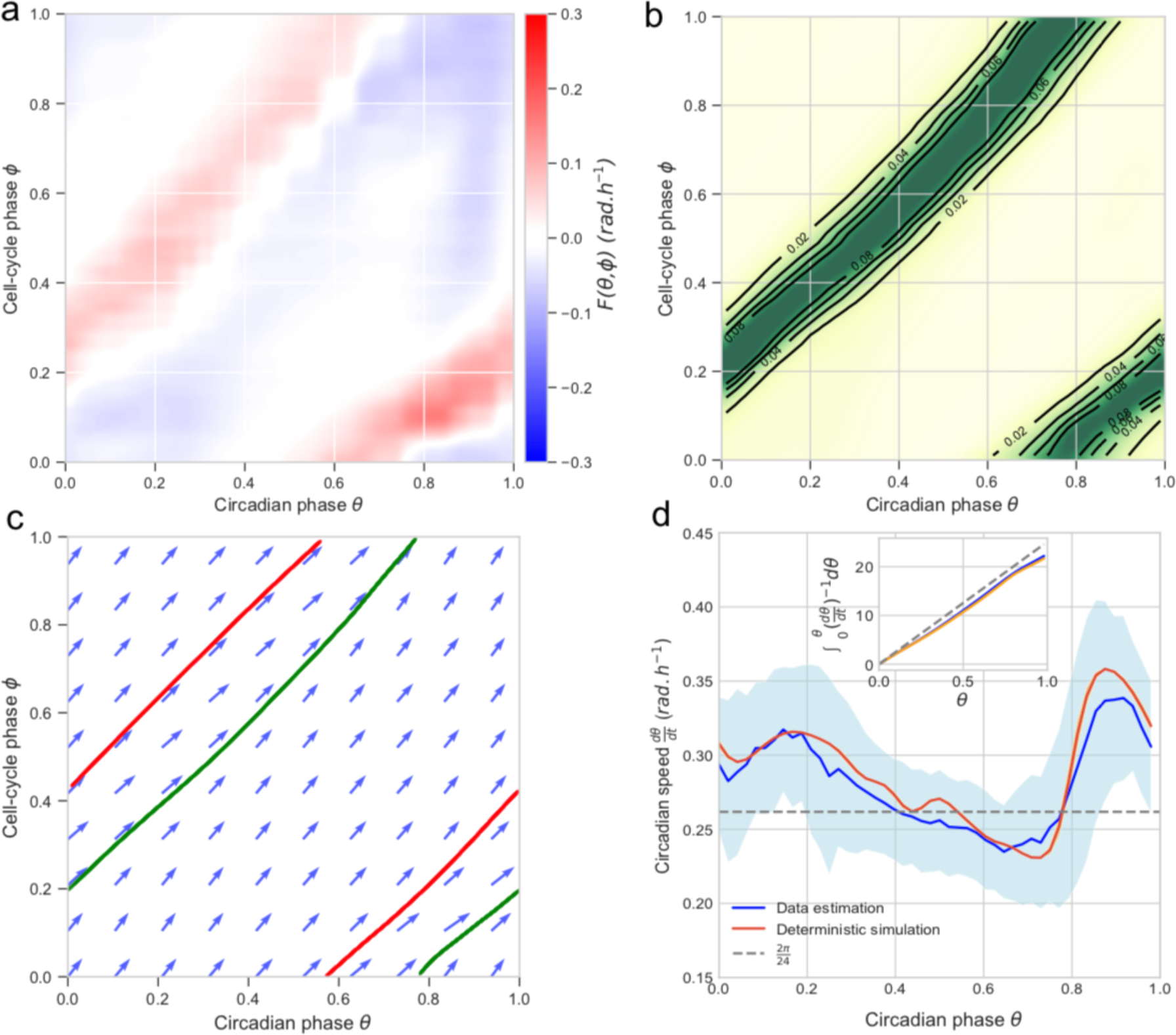
Influence of the cell cycle on the circadian phase enables 1:1 phase-locking. (**a**) Coupling *F*(*θ*, *φ*) optimized on dividing single-cell traces (all temperatures are pooled, cf. Supplementary Fig. 2). (**b**) Density of inferred phase traces from all the dividing traces with a 22±1h cell-cycle intervals indicates a 1:1 phase-locked state. (**c**) Numerical integration of phase velocity field (arrows, det erministic model) yields 1:1 attractor (green line) and repeller (red line). Here, the cell-cycle period was set to 22 h. (**d**) Circadian phase velocity speed is not constant, here for cells with 22±1h cell-cycle intervals. Data (blue line, standard deviation in light-blue shading) and deterministic simulation (orange line). Inset: integrated time along the attractor. The gray line shows constant bare phase velocity 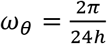.

The phase density for a fixed cell-cycle period of 22h (corresponding to the mean cell-cycle periods in the full dataset) (Fig. 2b, and Movie 1) clearly suggests 1:1 phase-locking. In fact, analyzing the predicted deterministic dynamics (Equation 1, with the reconstructed *F*(*θ*, *φ*), and without the noise) shows a 1:1 attractor (Fig. 2c). Thus, in this 1:1 state, the endogenous circadian period of 24h is shortened by two hours, which results from acceleration occurring after cytokinesis (*φ* = 0) when the circadian phase is near *θ* ≈ 0.8×2*π*, and lasting for the entire G1 phase, until about *θ* ≈ 0.4×2*π* when cells typically enter S phase (*φ* ≈ 0.4×2*π* at the G1/S transition, Supplementary Information) (Fig. 2d).

In sum, we reconstructed a coupling function from time-lapse cell traces that explains how the circadian phase velocity is adjusted throughout the cell cycle to produce 1:1 phase-locking.

### The inferred coupling predicts phase dynamics in perturbation experiments

The reconstructed model allows us to simulate the circadian phase dynamics in function of the cell-cycle period, which is relevant as the cells display a significant range of cell-cycle lengths (Supplementary Fig. 3a). In the deterministic system, we find 1:1 phase-locking over a range of cell-cycle times varying from 19h to 27h, showing that the cell cycle can both globally accelerate and slow down circadian phase progression (Fig. 3a). The attractor shifts progressively to the right in the phase-space, yielding a circadian phase at division ranging from *θ* ≈ 0.7×2*π* at division when *T*_*φ*_ = 19*h* to *θ* ≈ 0.9×2*π* when *T*_*φ*_ = 27*h*. Since the attractor for different cell-cycle periods shifts, the circadian phase velocity profile also changes (Supplementary Fig. 3b). To validate the predicted shifts, we experimentally subjected cells to perturbations inducing a large variety of cell-cycle periods, and compared the observed circadian phase to the model prediction at three different cell-cycle phases, revealing an excellent agreement, with no additional free parameters (Fig. 3b).

**Figure 3:**
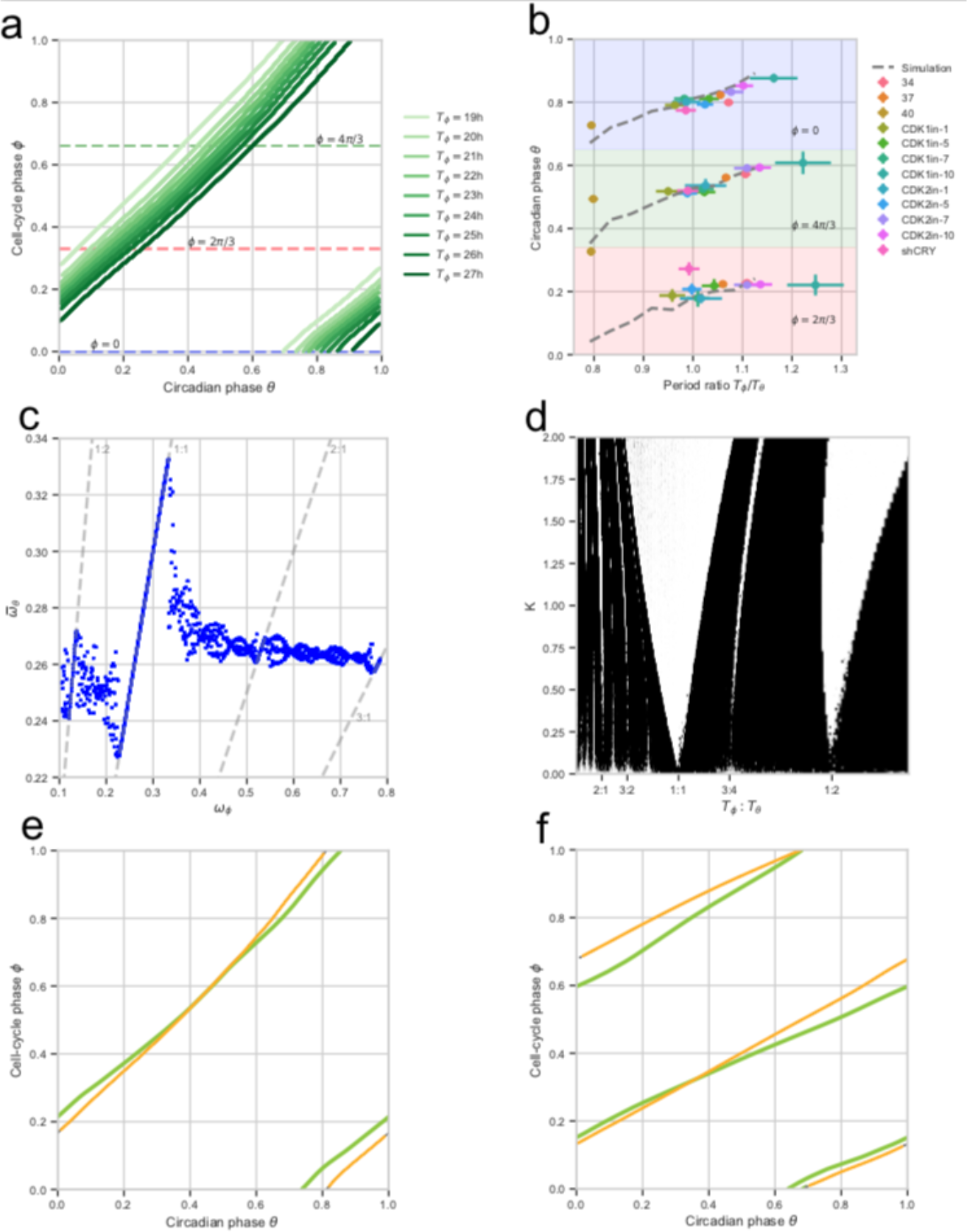
The coupling between the cell cycle and the circadian oscillator predicts phase shifts and phase-locking attractors in perturbation experiments. (**a**) Simulated (deterministic) attractors for cell-cycle periods ranging from 19h to 27h show that the dephasing of the cell cycle and the circadian oscillator changes within the 1:1 state. Periods just outside of this range yield quasiperiodic orbits. The horizontal dashed lines indicate three different cell-cycle phases *φ* = 0, 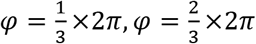 used in panel (b). (**b**) Predictions from a) (dashed grey lines) against independent experimental data collected from 12 perturbation experiments (colored symbols, see legend, notation explained in Methods). (**c**) Multiple phased-locked are predicted, recognizable by rational relationships between the frequencies of the entraining (cell cycle) and the entrained circadian oscillator, interspersed by quasiperiodic intervals. (**d**) Arnold tongues showing multiple phase-locked states in function of cell-cycle periods and coupling strength (*K*=1 corresponds to the experimentally found coupling). Stable zones (white tongues) reveal attractors interspersed by quasi-periodicity. Although there are only two wider phased-locked state (1:1 and 1:2), several other *p*:*q* phase-lockings are found. (**e-f**) Representative single-cell traces (data in yellow) evolving near predicted attractors (green lines). A cell with *T*_*φ*_ = 24*h* (e) and one with *T*_*φ*_ = 48*h* are near the 1:1 and 1:2 orbits, respectively.

The simulations also clearly revealed multiple phase-locked states (Arnold Tongues) (1:2, 1:1, 2:1, 3:1, etc.) (Fig. 3c, and Movie 2 for an animated phase-space representation). In the data, we identified cell trajectories following almost perfectly the deterministic attractors, both in the 1:1 and 1:2 phase-locking states (Fig. 3e and 3f, respectively); however, cells showing other *p:q* states were rarely observed.

### Fluctuations extend 1:1 phase-locking asymmetrically

To understand the differences between the simulated deterministic system and observed cell traces, we simulated the stochastic dynamics (Equation 1). We then compared measured data trajectories stratified by cell-cycle period (Fig. 4a) with deterministic (Fig. 4b) and stochastic simulations (Fig. 4c). This revealed that data agree better with stochastic than deterministic simulations, indicating that the phase fluctuations qualitatively change the phase portrait. One striking observation is the increased range of 1:1 phase-locking in the noisy system, however asymmetrically, since this occurs for shorter, but not for longer cell-cycle periods. Indeed, while 1:2 phase-locking is observed in the data and the noisy simulations, the deterministically predicted 2:1 state is replaced in the data and stochastic system by 1:1-like orbits. Consistently, spectral analysis revealed significant differences between deterministic and stochastic simulations (Supplementary Fig. 4a-b, Movie 3); in addition, the coupling, specifically in the 1:1 state, was able to efficiently filter the noise (Supplementary Fig. 4b, right).

**Figure 4:**
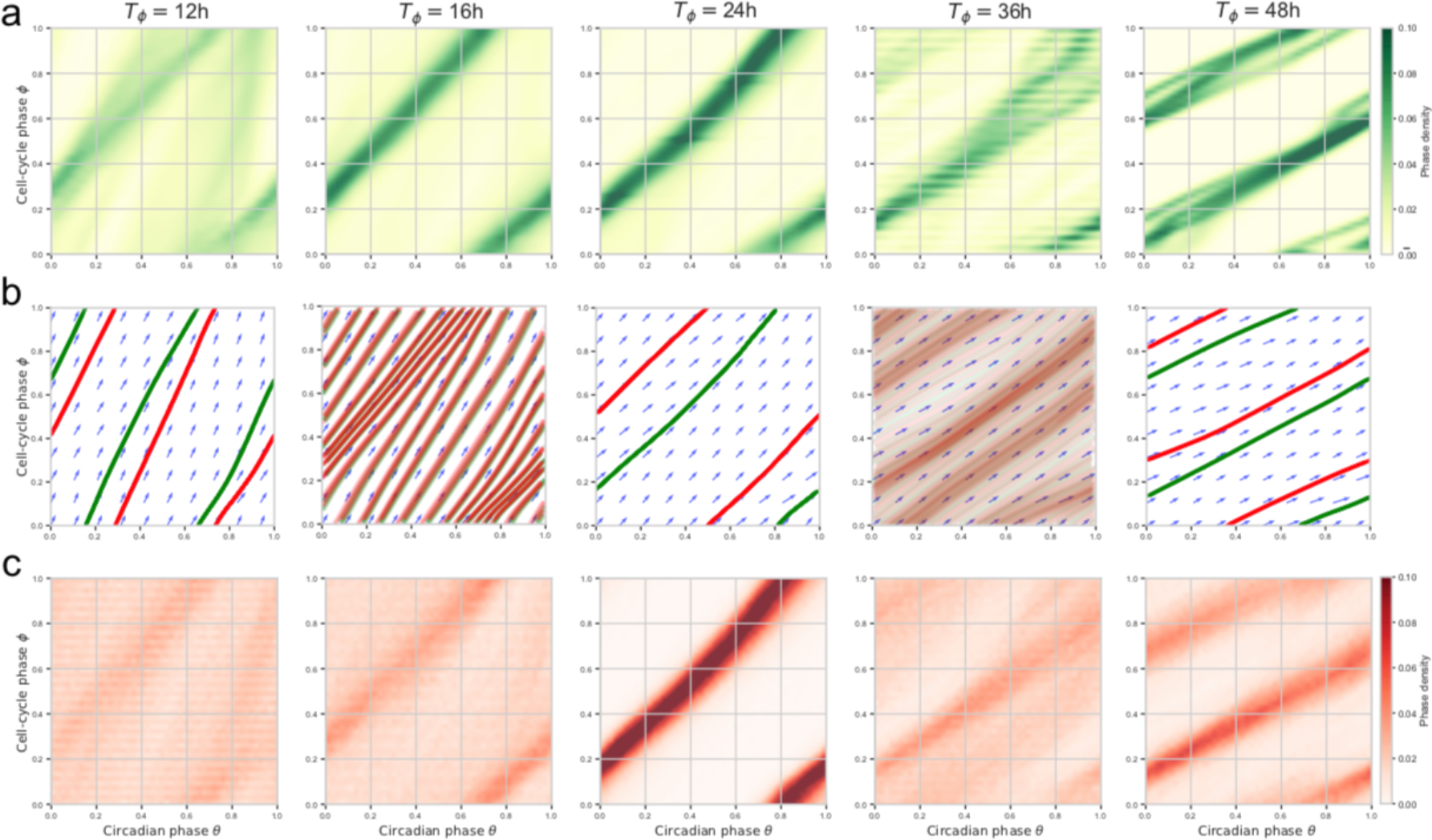
Single-cell data and stochastic simulations reveal robust 1:1 and 1:2 phase-locked states. (**a**) Phase-space densities of traces from the experimental traces stratified by cell-cycle period (±1h for each reference period). (**b**) Vector fields and simulated (deterministic) trajectories for the different cell-cycle period. Attractors are shown in green (forward time integration) and repellers (backward integration) in red (see also Movie 2). (**c**) Phase-space densities obtained from stochastic simulations of the model match better with the data compared to b).

### Evolutionarily conserved influence of the cell cycle on the circadian clock

Most studies on interacting cell cycle and circadian oscillators in mammals are from rodent models^1,5–7,9,20^. To test if the above dynamics are conserved in human U2OS cells, an established circadian oscillator model^21,22^, we engineered a U2OS cell line termed U2OS-Dual expressing a dual circadian fluorescent (Rev-Erb-α-YFP) and luminescent (Bmal1-luc) reporter system. U2OS-Dual cells possess a functional circadian clock behaving similarly to NIH3T3 cells also expressing a Bmal1-Luc luminescent reporter^23^ (Fig. 5a). To compare NIH3T3 and U2OS-Dual cells, we used our previous methods to segment, track and annotate traces obtained from 72h time-course under different conditions: at 34°C and at 37°C for cells with synchronized and non-synchronized circadian cycles^5^ (Fig. 5b).

**Figure 5:**
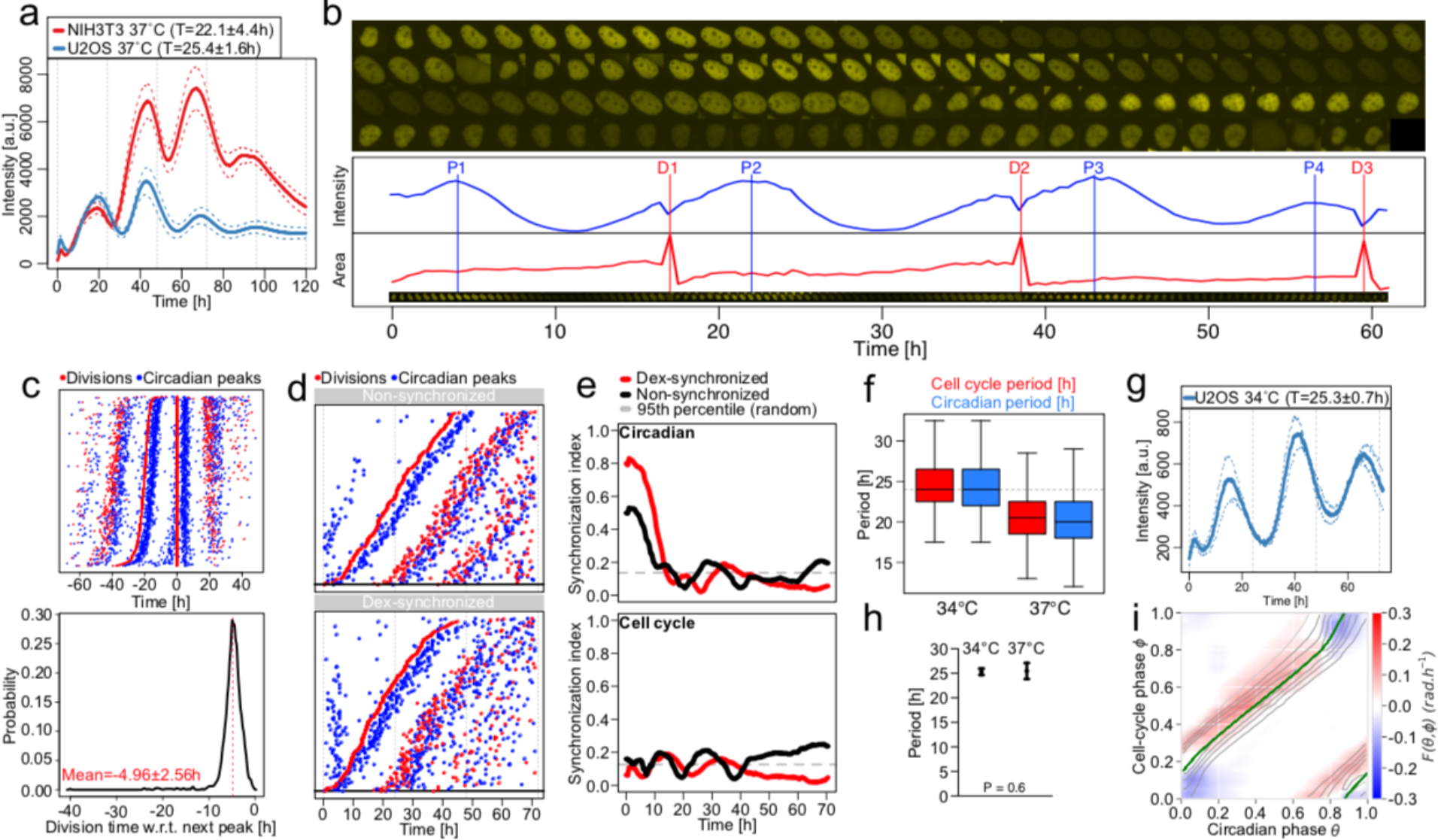
Conserved influence of the cell cycle on the circadian clock in human U2OS osteosarcoma cell. (**a**) Mean luminescence intensities (±SD, n=3) from non-dividing NIH3T3 and U2OS cells grown at 37°C expressing a Bmal1-Luc reporter. Values in the legend correspond to the mean periods ±SD. (**b**) Semi-automated segmentation and tracking of U2OS cell lines expressing the Rev-Erb-α-YFP circadian fluorescent reporter. Red vertical lines represent cell divisions (cytokinesis) and blue vertical lines show Rev-Erb-α-YFP signal peaks. (**c**) Top: stack of divisions (red) and Rev-Erb-α-YFP peaks (blue) for single U2OS traces centered on divisions. Bottom: distribution of the time of division relative to the next Rev-Erb-α-YFP peaks. (**d**) Divisions and Rev-Erb-α-YFP peaks from single non-synchronized (top), and dexamethasone(dex)-synchronized (bottom) U2OS traces ordered on the first division. (**e**) Synchronization index (SI) from non-synchronized (black) and dex-synchronized (red) traces for the circadian phase (top) and cell-cycle phase (bottom) estimated as in Bieler et al. ^5^. The circadian SI from non-synchronized cells is relatively high due to plating. Dashed gray lines show 95^th^ percentiles of the SI for randomly shuffled traces. (**f**) Cell-cycle and circadian periods for U2OS cells grown at 34°C and 37°C. Mean luminescence intensities (±SD, n=3) for non-dividing U2OS cells grown at 34°C expressing a Bmal1-Luc reporter. Values in the legend correspond to the mean periods ±SD. (**h**) Mean and standard deviation of the circadian period for non-dividing U2OS cells grown at 34°C and 37°C. (**i**) Coupling function *F*(*θ*, *φ*) optimized on all dividing U2OS traces grown at 37°C, superimposed with the attractor (*T*_*φ*_=22h) obtained from deterministic simulations (green line).

Similarly to NIH3T3 cells, the division events of non-synchronized U2OS-Dual cells grown at 37°C occurred 4.96 ± 2.6 h before a peak in the circadian fluorescent signal (Fig. 5c), indicating that the cell cycle and the circadian clock interact. To investigate the directionality of this interaction, we tested, like in NIH3T3 cells^5^, whether the circadian clock phase could influence cell-cycle progression by resetting the circadian oscillator using dexamethasone (dex), a well-established circadian resetting cue^24^ that does not perturb the cell-cycle^1^. We found the expected resetting effect of dex on the circadian phase by the density of peaks in reporter levels during the first 10h of recording, but with unnoticeable effects on the timing of the first division (Fig. 5d). However, the circadian peak following the first division occurred systematically around 5h after the division in both conditions, suggesting that cell division in U2OS can reset circadian phases and overwrite dex synchronization. Synchronization of the circadian clocks for dex-*vs* non-treated cells was expectedly higher for dex and gradually decreased to reach the level of the untreated cells (Fig. 5e), contrasting with the generally lower synchronization of cell divisions in both conditions. To then test if the cell cycle could influence the circadian clock, we lengthened the cell-cycle period by growing cells at 34°C and compared with 37°C. Interestingly, cells at 34°C showed a longer circadian period compared to 37°C (Fig. 5f), unlike the temperature compensated circadian periods (∼25h) in non-dividing cells (Fig. 5a, g-h).

Thus, similarly to mouse NIH3T3 cells, the coupling directionality is predominantly from the cell cycle to the circadian clock. In fact, the reconstructed coupling function for U2OS-Dual cells grown at 37°C (Fig. 5i) is structurally similar to that obtained in mouse fibroblasts (Fig. 2a), showing a 1:1 attractor. Taken together, these analyses revealed that the interaction between the cell cycle and the circadian clock is conserved in mouse and human cells.

### Circadian oscillations in dividing cells lose entrainment to temperature cycles

In mammals, circadian clocks in tissues are synchronized by multiple systemic signals^25^. In fact, temperature oscillations mimicking those physiologically observed can phase-lock circadian oscillators in non-dividing (contact-inhibited) NIH3T3 cells *in vitro*^26^. To study how the found interactions influence temperature entrainment, we applied temperature cycles (24h period ranging from 35.5°C to 38.5°C) to U2OS cells growing at different rates (plated at different densities) and monitored population-wide Bmal1-luc signals (Fig. 6a). Since temperature impacts the enzymatic activity of the luciferase^27^, we first corrected the signal for this systematic effect (Supplementary Information). We found that, independently of initial densities, as the populations reach confluency (which takes longer for initial low density), the phases and amplitudes become stationary, showing 1:1 entrainment (Fig. 6b, c and Supplementary Fig. 5a). During the initial transients, emerging circadian oscillations in non-confluent cells showed phases that were already stationary, at least once cell numbers were sufficiently high to obtain reliable signals.

**Figure 6:**
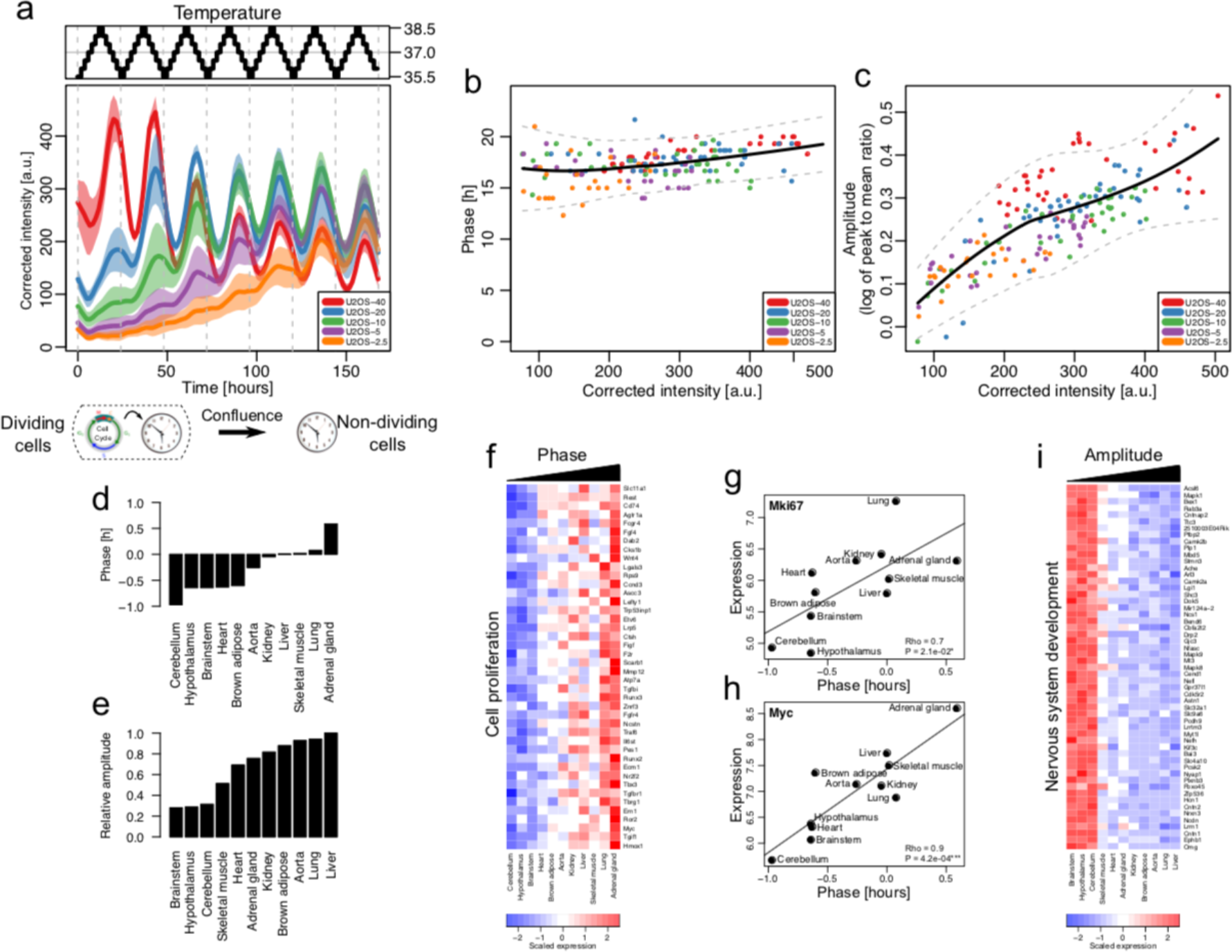
Temperature cycles do not entrain circadian oscillators in dividing cells and proliferation genes explains tissue-specific circadian phases. (**a**) Corrected and averaged Bmal1-Luc intensities and 95% confidence intervals (n=6) from U2OS-Dual cells plated at different initial densities and subjected to a temperature entrainment (top). (**b**) Acrophases of the Bmal1-Luc signal in function of the reporter intensity for the cells in a). Loess fit (black) and 95% confidence intervals (gray). (**c**) Amplitude (log of peak to mean ratio) of the Bmal1-Luc oscillations in function of the reporter intensity for the cells in a). Loess fit (black) and 95% confidence intervals (gray). (**d-e**) Circadian phases (d) and amplitudes (e) of different mouse tissues obtained in ref^29^, relative to liver. (**f**) Expression levels of genes positively associated with phases from d) and linked to cell proliferation. (**g-h**) Correlations between Mki67(g) and Myc(h) mRNA expression and circadian phases across mouse tissues. (**i**) Expression levels of genes negatively associated with amplitudes and linked to nervous system development.

As cell confluence increases, the proportions of cells which stop cycling (exit to G0) increases^28^. We therefore hypothesized that the observed phase and amplitude profiles in Bmal1-luc signals (Fig. 6b, c) originate from a mixture of two populations: an increasing population of non-dividing cells (G0) showing ‘normal’ entrainment properties, and dividing cells. We considered three scenarios for the dividing cells: i) the circadian oscillators in dividing cells adopt the same circadian profile as non-dividing entrained cells; ii) are not entrained; or iii) are entrained, but with a different phase compared to non-dividing cells (Supplementary Fig. 5b). These scenarios can be distinguished by the predicted phase and amplitude profiles (Supplementary Fig. 5c). Clearly, the measured profiles for U2OS-Dual cells favored the second scenario, suggesting that circadian oscillators in dividing cells do not entrain to the applied temperature cycles.

### Expression levels of proliferation genes explain tissue-specific circadian phases

Our findings suggest that phases or amplitudes of circadian clocks in organs *in vivo* might be influenced by the proliferation state of cells in the tissue. To test this, we first studied which cellular processes correlate with circadian clock parameters in different mouse tissues. A study of mRNA levels in twelve adult (6 weeks old males) mouse tissues revealed that clock phases span 1.5 hours between the earliest and latest tissues (Fig. 6d)^29,30^, an effect which is considered large in chronobiology since even period phenotypes of core clock genes are often smaller^2,31,32^. Using those data, we noticed that the mean mRNA levels across tissues of many genes correlated with the phase offsets (Supplementary Table 1). However, gene functions related to cell proliferation stood out as the most strongly enriched (Fig. 6f, Supplementary Table 1). Among those genes, the levels of known markers of cell proliferation such as *Mki67* or *Myc* were strongly correlated with the phase offsets (Fig. 6g-h, Supplementary Table 1). Amplitudes, on the other hand, were not correlated with proliferation genes, but rather with neuronal specific genes, as expected owing to the damped rhythms present in those tissues (Fig. 6i, Supplementary Table 1)^30^. Thus, even basal level of proliferation observed in normal tissues could explain the dephasing of the circadian clock, hence suggesting a physiological role for the described interaction of the cell cycle and circadian clocks.

## Discussion

A goal in quantitative single-cell biology is to obtain data-driven and dynamical models of biological phenomena in low dimensions. In practice, the heterogeneity and complex physics underlying the emergence of structure and function in non-equilibrium living systems, as well as the sparseness of available measurements pose challenges. Here, we studied a system of two coupled biological oscillators, sufficiently simple to allow data-driven model identification, yet complex enough to exhibit qualitatively distinct dynamics, *i.e. p:q* states and quasi-periodicity. This revealed that the coupling of cell cycle and circadian oscillators only depends weakly on temperature, and is also conserved from mouse to human cells. The coupling predicted multiple phase-locked states that showed different levels of stability against fluctuations. In a physiological context, the cell proliferation status in mouse tissues explained tissue-specific circadian phases.

In the coupled cell cycle and circadian oscillator system, phase-locked states different from 1:1 have been observed, despite the fact that oscillator theory^33^, as well as specific models of this system in cyanobacteria^34^, indicate that noise tends to destabilize higher order phase-locking. In single cyanobacteria cells, a transition from 1:1 to 2:1 was found when growth rates were increased under constant light, which was modeled as caused by the unilateral and localized influence of the circadian clock on the cell-cycle progression, such that the 2:1 state corresponded to two cell divisions every 24h^11^. While multiple attractors, notably 1:1 and 3:2, were observed in NIH3T3 cells under transient dexamethasone stimulation^6^, we here report 1:2 states for long cell-cycle times under steady, unstimulated, conditions. Unlike other deterministically predicted state, such as 2:1, which were less stable against noise and rarely observed, 1:2 was sufficiently robust and observed in cells.

In fact, we found that noise extended the range of our 1:1 tongue, but asymmetrically, *i.e.* towards decreased cell-cycle periods. This may be reminiscent of generalized Poincaré oscillators showing that the entrainment range is broader for limit cycles with low relaxation rates^35^. Indeed, noise could decrease relaxation rates and thereby broaden Arnold tongues. While other works have explained the phase-locked states either fully or partially due to the influence of the circadian oscillator on cell-cycle progression^6,7,11^, the mammalian cells analyzed here showed that the locked states follow only from the influence of the cell cycle on the circadian phase progression, as represented through the reconstructed function *F*(*θ*, *φ*), both in mouse and human cells.

In mammals, the circadian oscillator in the suprachiasmatic nucleus (SCN) is the pacemaker for the entire organism^36^, driving 24h rhythms in activity, feeding, body temperature and hormone levels. In particular, the SCN can synchronize peripheral cell-autonomous circadian clocks located within organs across the body^37^. Consistent with theory^38^, in a physiological context of entrainment, the coupling of the cell cycle with the circadian clock can induce proliferation dependent phase-shifts, which we observed. Such phase shifts could reflect a homogenous behavior of all cells, or it could reflect an underlying heterogeneity of cell proliferation states, for example in organs with a complex substructure such as the liver^39^, where parenchymal cells might have different turnover rates compared to other cells. Such situations might then lead to local phase heterogeneity and possibly wave propagation. The phase shifts we observed in tissues were associated with low proliferation, *i.e.* non-pathological states of tissue homeostasis and cell renewal. For example, the liver or the adrenal gland showed a phase advance compared to fully quiescent tissues like the brain.

When cell proliferation is abnormally high such as in cancer, circadian clocks are often severely damped^40^. While this absence of a robust circadian rhythm in malignant tissue states may reflect non functional circadian oscillators due to mutations in clock genes^41^, the damped rhythms may also reflect circadian desynchrony of otherwise functional circadian oscillators. Such desynchrony would readily follow from the coupling between the cell-cycle and circadian oscillator we highlight here, in the presence of non-coherent cell cycle progression.

Methodologically, the new approach to reconstruct a dynamical model for the coupled oscillator system has significant advantages over previous methods, notably strong assumptions such as the sparse and localized coupling^5^ are dispensable. This revealed a coupling function composed of diffuse regions of phase acceleration and deceleration leading to robust 1:1 phase locking. Compared with generic model identification techniques^42^, our approach models the raw data and its noise structure explicitly. In the future, such data-driven identification of dynamical models might reveal dynamical instabilities underlying ordered states in spatially extended systems, as occurring, for instance, during somitogenesis^43^.

## Methods

### Cell lines

All cell lines (U2OS-Dual, NIH3T3-Bmal1-Luc, and U2OS-PGK-Luc) were maintained in a humidified incubator at 37°C with 5% CO2 using DMEM cell culture media supplemented with 10% fetal bovine serum (FBS) and 1% penicillin-streptomycin-glutamine (PSG). One day before luminescence or fluorescence acquisitions, we replaced DMEM with FluoroBrite DMEM media supplemented with 10% FBS and 1% PSG. NIH3T3 perturbation experiments were generated in Bieler et al^5^. Briefly, they correspond to temperature changes (34, 37, and 40°C), treatment with CDK1 (RO-3306, Sigma-Aldrich) and CDK1/2 (NU-6102, Calbiochem) inhibitors at 1, 5, 7 and 10μM (CDK1in-[1,5,7,10] and CDK2in-[1,5,7,10]), and shRNA-mediated knockdown of Cry2 (shCry).

### Fluorescent time-lapse microscopy

Time-lapse fluorescent microscopy for U2OS-Dual cells was performed at the biomolecular screening facility (BSF, EPFL) using an InCell Analyzer 2200 (GE healthcare). Experiments were performed at different temperature (34°C, 37°C, or 40°C) with a humidity and CO_2_ (5%) control system. We used 100 ms excitation at 513/17 nm and emission at 548/22 nm to record the YFP channel. Cells were recorded by acquiring one field of view per well in a 96-well black plate (GE healthcare). We used our previously developed semi-automated pipeline for segmentation and tracking of individual cells^5^.

### Probabilistic model for reconstructing phase dynamics and coupling of two biological oscillators

Denote by ***D*** the entire set of single cells comprising temporal intensity measurements (Δ*t*=30 min) from all fluorescent traces and **Λ** the set of model parameters, comprising the gridded coupling function *F*_*ij*_. To reconstruct the phase dynamics of our model, we seek to maximize the likelihood of the data 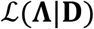 that is, we solve:

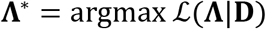

In practice, we used an EM algorithm, by iteratively optimizing the function *Q*(**Λ**, **Λ**′) over its first argument, where *Q* can be written as follow:

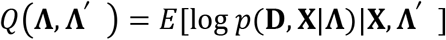

That is, *Q*(**Λ**, **Λ**′) corresponds to the expected value of the log-likelihood of the data with respect to the posterior probabilities of the hidden phases **X** (latent variables), computed using the current parameter **Λ**′. This process guarantees a monotonous convergence of the log-likelihood, although a global maximum is not necessarily reached^44^.

To control for the many parameters *F*_*ij*_, we added regularization constraints for both the smoothness and sparsity:

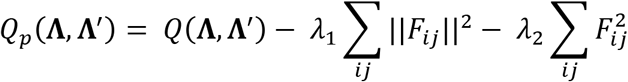

This expression is also guaranteed to converge^45^.

Details about the optimization method, choice of the regularization parameters, and computation of the phase posteriors using a HMM are provided in Supplementary Information.

### Long-term temperature entrainment and luminescence recording

We performed long-term temperature entrainment experiments using a Tecan plate reader Infinite F200 pro with a CO_2_ and temperature modules. One day before starting the experiment, serial dilution ranging from 40,000 to 2,500 cells were seeded in 96-well white flat bottom plates (Costar 3917). To prevent media evaporation, all wells were filled with 300μl of media composed of FluoroBrite, 10% FBS, 1% PSG, and 100 nM D-luciferin (NanoLight technology) and covered with a sealing tape (Costar 6524). We set up temperature entrainment using stepwise increase (or decrease) of 0.5°C every 2 hours to produce temperature oscillating profiles going from 35.5°C to 38.5°C and back to 35.5°C again over a period of 24 hours. Intensities from all wells were recorded every 10 minutes with an integration time of 5000 milliseconds.

### Testing the association between averaged gene expression and phases or amplitudes in different mouse tissues

We used the average gene expression obtained from a selected set of twelve adult (6 weeks old males) mouse tissues from the Zhang et al. dataset (GEO accession GSE54650)^30^. For this analysis, we estimated the Pearson’s correlation between the averaged gene expression and the circadian tissue phases or amplitudes reported in ref^29^. We selected the top 200 genes positively or negatively associated with either the phases or the amplitudes for gene ontology analysis^46^ (Supplementary Table 1).

## Data availability

The data supporting figures and other findings of this study are available from the corresponding author on request.

## Code availability

The code is available online at the following URL: https://github.com/ColasDroin/CouplingHMM.

## Acknowledgments

We thank Rosamaria Cannavo for engineering the U2OS-Dual cell line and Jonathan Bieler for initial analyses. Fabien Kuttler from the Swiss Federal Institute of Technology (EPFL) Biomolecular Screening Facility (EPFL-BSF) and Luigi Bozzo and José Artacho from the EPFL Bioimaging and Optics Core Facility (EPFL-BIOP) for assistance with the imaging. Fluorescence-activated cell sorting was performed at the EPFL Flow Cytometry Core Facility (EPFL-FCCF). This work was supported by the Swiss National Science Foundation Grant 310030_173079 and the EPFL. E.R.P was supported by a Canadian Institute of Health Research (CIHR 358808) and a SystemsX.ch Transition Postdoc Fellowships (51FSP0163584).

## Contributions

C.D., E.R.P., and F.N. designed and participated in the study concept. C.D. and E.R.P. developed computational analysis tools. E.R.P. performed the experiments. C.D. and E.R.P. processed and analyzed the experimental data. C.D., E.R.P., and F.N. interpreted the results.

E.R.P. and F.N. acquired the funding. F.N. supervised the study. C.D., E.R.P., and F.N. wrote the manuscript.

---

**Movie 1: Single-cell trajectories of circadian and cell-cycle phases cluster around a 1:1 phase-locked state.**

Inferred temporal single-cell trajectories in the phase-space. Each cell is represented by a point, and its instantaneous direction by a tail. The color shows the instantaneous phase velocity of the circadian phase of the cell is blue or red depending in the instantaneous speed is respectively slower of faster than the intrinsic circadian clock speed 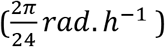. The phase density is progressively built in the background from the accumulation of the cell trajectories.

**Movie 2: Simulations of the deterministic model shows phase-locking and quasi-periodicity**

Vector fields and simulated (deterministic) trajectories for increasing cell-cycle periods *T*_*φ*_. Simulations in forward time are represented with a green line, while simulations in backward time are represented with a red. The vector field (in blue) shows the instantaneous phase velocities. Phase-locked states, interspersed with quasi-periodic orbits, are successively observed. The beginnings of trajectories are removed to show steady-states.

**Movie 3: Spectral analysis of the deterministic and stochastic simulations shows qualitative differences in function of the cell-cycle period.**

Animated representation of Supplementary Figure 4 for increasing cell-cycle velocity. The instantaneous position of the system on the Arnold tongues is represented by a red dot.

## References

1 Nagoshi, E. et al. Circadian gene expression in individual fibroblasts: cell-autonomous and self-sustained oscillators pass time to daughter cells. Cell 119, 693–705, doi:10.1016/j.cell.2004.11.015 (2004).

2 Mermet, J., Yeung, J. & Naef, F. Systems Chronobiology: Global Analysis of Gene Regulation in a 24-Hour Periodic World. Cold Spring Harbor perspectives in biology 9, doi:10.1101/cshperspect.a028720 (2017).

3 Hahn, A. T., Jones, J. T. & Meyer, T. Quantitative analysis of cell cycle phase durations and PC12 differentiation using fluorescent biosensors. Cell cycle 8, 1044–1052, doi:10.4161/cc.8.7.8042 (2009).

4 Spencer, S. L. et al. The proliferation-quiescence decision is controlled by a bifurcation in CDK2 activity at mitotic exit. Cell 155, 369–383, doi:10.1016/j.cell.2013.08.062 (2013).

5 Bieler, J. et al. Robust synchronization of coupled circadian and cell cycle oscillators in single mammalian cells. Molecular systems biology 10, 739, doi:10.15252/msb.20145218 (2014).

6 Feillet, C. et al. Phase locking and multiple oscillating attractors for the coupled mammalian clock and cell cycle. Proceedings of the National Academy of Sciences of the United States of America 111, 9828–9833, doi:10.1073/pnas.1320474111 (2014).

7 Matsuo, T. et al. Control mechanism of the circadian clock for timing of cell division in vivo. Science 302, 255–259, doi:10.1126/science.1086271 (2003).

8 Glass, L. Synchronization and rhythmic processes in physiology. Nature 410, 277–284, doi:10.1038/35065745 (2001).

9 Kowalska, E. et al. NONO couples the circadian clock to the cell cycle. Proceedings of the National Academy of Sciences of the United States of America 110, 1592–1599, doi:10.1073/pnas.1213317110 (2013).

10 Mori, T., Binder, B. & Johnson, C. H. Circadian gating of cell division in cyanobacteria growing with average doubling times of less than 24 hours. Proceedings of the National Academy of Sciences of the United States of America 93, 10183–10188 (1996).

11 Yang, Q., Pando, B. F., Dong, G., Golden, S. S. & van Oudenaarden, A. Circadian gating of the cell cycle revealed in single cyanobacterial cells. Science 327, 1522–1526, doi:10.1126/science.1181759 (2010).

12 Matsu-Ura, T. et al. Intercellular Coupling of the Cell Cycle and Circadian Clock in Adult Stem Cell Culture. Molecular cell 64, 900–912, doi:10.1016/j.molcel.2016.10.015 (2016).

13 Plikus, M. V. et al. Local circadian clock gates cell cycle progression of transient amplifying cells during regenerative hair cycling. Proceedings of the National Academy of Sciences of the United States of America 110, E2106–2115, doi:10.1073/pnas.1215935110 (2013).

14 Paijmans, J., Bosman, M., Ten Wolde, P. R. & Lubensky, D. K. Discrete gene replication events drive coupling between the cell cycle and circadian clocks. Proceedings of the National Academy of Sciences of the United States of America 113, 4063–4068, doi:10.1073/pnas.1507291113 (2016).

15 Shostak, A. et al. MYC/MIZ1-dependent gene repression inversely coordinates the circadian clock with cell cycle and proliferation. Nature communications 7, 11807, doi:10.1038/ncomms11807 (2016).

16 Rougemont, J. & Naef, F. Collective synchronization in populations of globally coupled phase oscillators with drifting frequencies. Physical review. E, Statistical, nonlinear, and soft matter physics 73, 011104, doi:10.1103/PhysRevE.73.011104 (2006).

17 d’Eysmond, T., De Simone, A. & Naef, F. Analysis of precision in chemical oscillators: implications for circadian clocks. Phys Biol 10, 056005, doi:10.1088/1478-3975/10/5/056005 (2013).

18 Teramae, J. N., Nakao, H. & Ermentrout, G. B. Stochastic phase reduction for a general class of noisy limit cycle oscillators. Physical review letters 102, 194102, doi:10.1103/PhysRevLett.102.194102 (2009).

19 Rabiner, L. R. & Juang, B.-H. An introduction to hidden Markov models. ieee assp magazine 3, 4–16 (1986).

20 Yeom, M., Pendergast, J. S., Ohmiya, Y. & Yamazaki, S. Circadian-independent cell mitosis in immortalized fibroblasts. Proceedings of the National Academy of Sciences of the United States of America 107, 9665–9670, doi:10.1073/pnas.0914078107 (2010).

21 Vollmers, C., Panda, S. & DiTacchio, L. A high-throughput assay for siRNA-based circadian screens in human U2OS cells. PloS one 3, e3457, doi:10.1371/journal.pone.0003457 (2008).

22 Maier, B. et al. A large-scale functional RNAi screen reveals a role for CK2 in the mammalian circadian clock. Genes & development 23, 708–718, doi:10.1101/gad.512209 (2009).

23 Nicolas, D., Zoller, B., Suter, D. M. & Naef, F. Modulation of transcriptional burst frequency by histone acetylation. Proceedings of the National Academy of Sciences of the United States of America 115, 7153–7158, doi:10.1073/pnas.1722330115 (2018).

24 Balsalobre, A. et al. Resetting of circadian time in peripheral tissues by glucocorticoid signaling. Science 289, 2344–2347 (2000).

25 Dibner, C., Schibler, U. & Albrecht, U. The mammalian circadian timing system: organization and coordination of central and peripheral clocks. Annual review of physiology 72, 517–549, doi:10.1146/annurev-physiol-021909-135821 (2010).

26 Saini, C., Morf, J., Stratmann, M., Gos, P. & Schibler, U. Simulated body temperature rhythms reveal the phase-shifting behavior and plasticity of mammalian circadian oscillators. Genes & development 26, 567–580, doi:10.1101/gad.183251.111 (2012).

27 Koksharov, M. I. & Ugarova, N. N. Approaches to engineer stability of beetle luciferases. Computational and structural biotechnology journal 2, e201209004, doi:10.5936/csbj.201209004 (2012).

28 Hayes, O. et al. Cell confluency is as efficient as serum starvation for inducing arrest in the G0/G1 phase of the cell cycle in granulosa and fibroblast cells of cattle. Animal reproduction science 87, 181–192, doi:10.1016/j.anireprosci.2004.11.011 (2005).

29 Yeung, J. et al. Transcription factor activity rhythms and tissue-specific chromatin interactions explain circadian gene expression across organs. Genome research 28, 182–191, doi:10.1101/gr.222430.117 (2018).

30 Zhang, R., Lahens, N. F., Ballance, H. I., Hughes, M. E. & Hogenesch, J. B. A circadian gene expression atlas in mammals: implications for biology and medicine. Proceedings of the National Academy of Sciences of the United States of America 111, 16219–16224, doi:10.1073/pnas.1408886111 (2014).

31 Cermakian, N., Monaco, L., Pando, M. P., Dierich, A. & Sassone-Corsi, P. Altered behavioral rhythms and clock gene expression in mice with a targeted mutation in the Period1 gene. The EMBO journal 20, 3967–3974, doi:10.1093/emboj/20.15.3967 (2001).

32 Debruyne, J. P. et al. A clock shock: mouse CLOCK is not required for circadian oscillator function. Neuron 50, 465–477, doi:10.1016/j.neuron.2006.03.041 (2006).

33 Guckenheimer, J. & Holmes, P. Nonlinear oscillations, dynamical systems, and bifurcations of vector fields. Vol. 42 %@ 1461211409 (Springer Science & Business Media, 2013).

34 Monti, M., Lubensky, D. K. & Ten Wolde, P. R. Optimal entrainment of circadian clocks in the presence of noise. Physical review. E 97, 032405, doi:10.1103/PhysRevE.97.032405 (2018).

35 Granada, A. E. & Herzel, H. How to achieve fast entrainment? The timescale to synchronization. PloS one 4, e7057, doi:10.1371/journal.pone.0007057 (2009).

36 Hastings, M. H., Maywood, E. S. & Brancaccio, M. Generation of circadian rhythms in the suprachiasmatic nucleus. Nature reviews. Neuroscience 19, 453–469, doi:10.1038/s41583-018-0026-z (2018).

37 Mohawk, J. A., Green, C. B. & Takahashi, J. S. Central and peripheral circadian clocks in mammals. Annual review of neuroscience 35, 445–462, doi:10.1146/annurev-neuro-060909-153128 (2012).

38 Pikovsky, A. R. M. K. J. Synchronization. (Cambridge University Press, 2001).

39 Halpern, K. B. et al. Single-cell spatial reconstruction reveals global division of labour in the mammalian liver. Nature 542, 352–356, doi:10.1038/nature21065 (2017).

40 Shilts, J., Chen, G. & Hughey, J. J. Evidence for widespread dysregulation of circadian clock progression in human cancer. PeerJ 6, e4327, doi:10.7717/peerj.4327 (2018).

41 Kelleher, F. C., Rao, A. & Maguire, A. Circadian molecular clocks and cancer. Cancer letters 342, 9–18, doi:10.1016/j.canlet.2013.09.040 (2014).

42 Brunton, S. L., Proctor, J. L. & Kutz, J. N. Discovering governing equations from data by sparse identification of nonlinear dynamical systems. Proceedings of the National Academy of Sciences of the United States of America 113, 3932–3937, doi:10.1073/pnas.1517384113 (2016).

43 Soroldoni, D. et al. Genetic oscillations. A Doppler effect in embryonic pattern formation. Science 345, 222–225, doi:10.1126/science.1253089 (2014).

44 Dempster, A. P., Laird, N. M. & Rubin, D. B. Maximum likelihood from incomplete data via the EM algorithm. Journal of the royal statistical society. Series B (methodological), 1–38 %@ 0035-9246 (1977).

45 Green, P. J. On use of the EM for penalized likelihood estimation. Journal of the Royal Statistical Society. Series B (Methodological), 443–452 %@ 0035-9246 (1990).

46 Huang da, W., Sherman, B. T. & Lempicki, R. A. Bioinformatics enrichment tools: paths toward the comprehensive functional analysis of large gene lists. Nucleic acids research 37, 1–13, doi:10.1093/nar/gkn923 (2009).

