## Supplementary figures for "Low-dimensional Dynamics of Two Coupled Biological Oscillators"

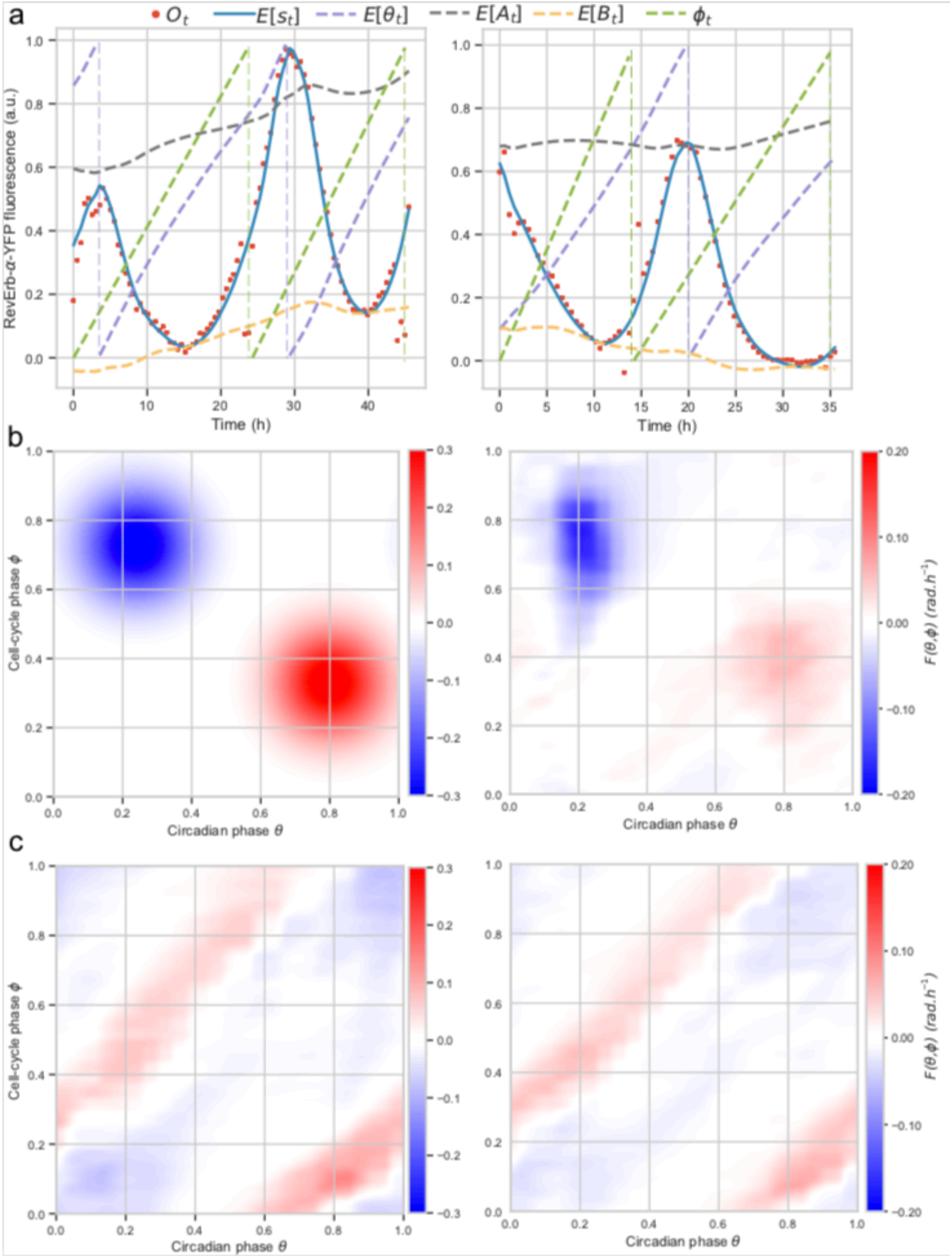


**Supplementary Figure 1: Validation of the inference methods.** (**a**) Examples of temporal data traces and model fits. The colors show data (red), posterior mean for $S_t$ (blue), posteriors circular mean for $\theta_t$ (dashed purple), posterior means $A_t$ (dashed grey) and baseline $B_t$ (dashed yellow). The cell-cycle phase $\varphi_t$ (which is not a hidden variable) is obtained from linear interpolation between two successive divisions (dashed green line). Deviations of the purple curve from a straight line corresponds to transient variations of circadian phase velocity, owing to noise and coupling. (**b-c**) Using oscillator parameters mimicking real cells, we can recover coupling functions from simulated traces. (**b**) First, we simulated traces with a coupling $F(\theta, \varphi)$ comprising two Gaussian interaction regions, as shown in the left panel. The reconstructed function is shown in the right panel. (**c**) Same numerical experiment made with the coupling inferred from the real data as input (left). Both simulations reveal that the inferred functions are qualitatively accurate, but quantitatively damped.

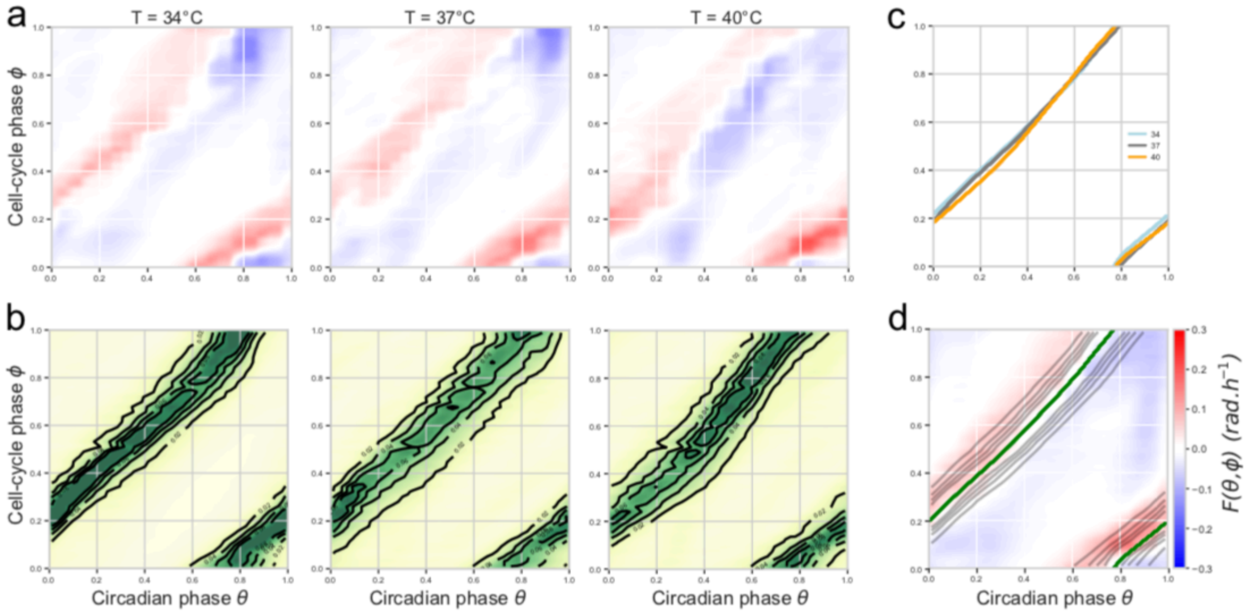


**Supplementary Figure 2: $F(\theta,\varphi)$ depends weakly on temperature.** (**a**) Coupling functions obtained from the traces acquired at 34°C (left), 37°C (middle) and 40°C (right). To avoid possible bias, the traces were sampled such that the distribution of cell-cycle periods is identical for each temperature (Supplementary Fig. 3a for the cell-cycle period distributions at the three different temperatures). (**b**) Superimposition of the merged (from all traces at 34°C, 37°C, and 40°C) coupling function (that of Fig. 2a), the phase-space trace density (Fig. 2b, here shown as contour lines), and the attractor (Fig. 2c, green line). (**c**) Attractors for the vector fields in a) show that the 1:1 phase-locked orbit is temperature independent. (**d**) Phase space densities obtained in the same condition and from the same traces as in (a).

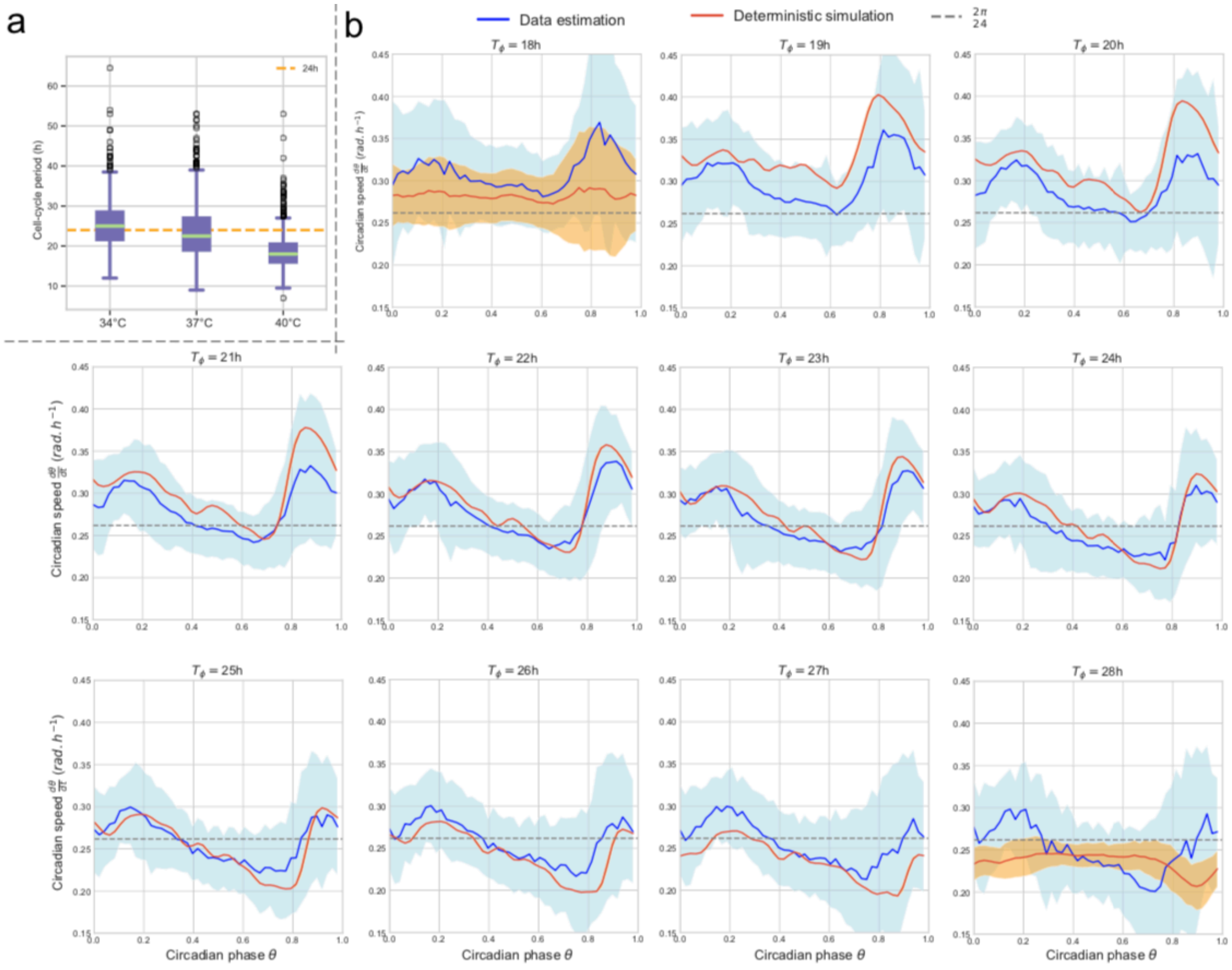


**Supplementary Figure 3: The phase velocity profiles along the 1:1 attractors in function of cell-cycle period.** (**a**) Cell-cycle period (division-to-division time intervals) distributions in NIH3T3 cells at three temperatures. The average period progressively decreases from around 24h at 34°C to 18h at 40°C[5]. (**b**) Circadian phase velocity on the 1:1 attracting orbit in function of cell-cycle period (increasing from left to right and top to bottom). The phase velocity shown is either inferred from the data traces (blue line, standard deviation in light blue), or simulated using the deterministic model (no phase noise) (orange line). The first and last panels ($T_{\varphi}$ = 18h and 28h) have quasi-periodic orbits (hence a standard deviation is associated with the mean phase velocity). The natural (non-dividing cells) circadian phase velocity (about 0.26 rad.h$^{-1}$, corresponding to a 24h period) is indicated by a dashed grey line.

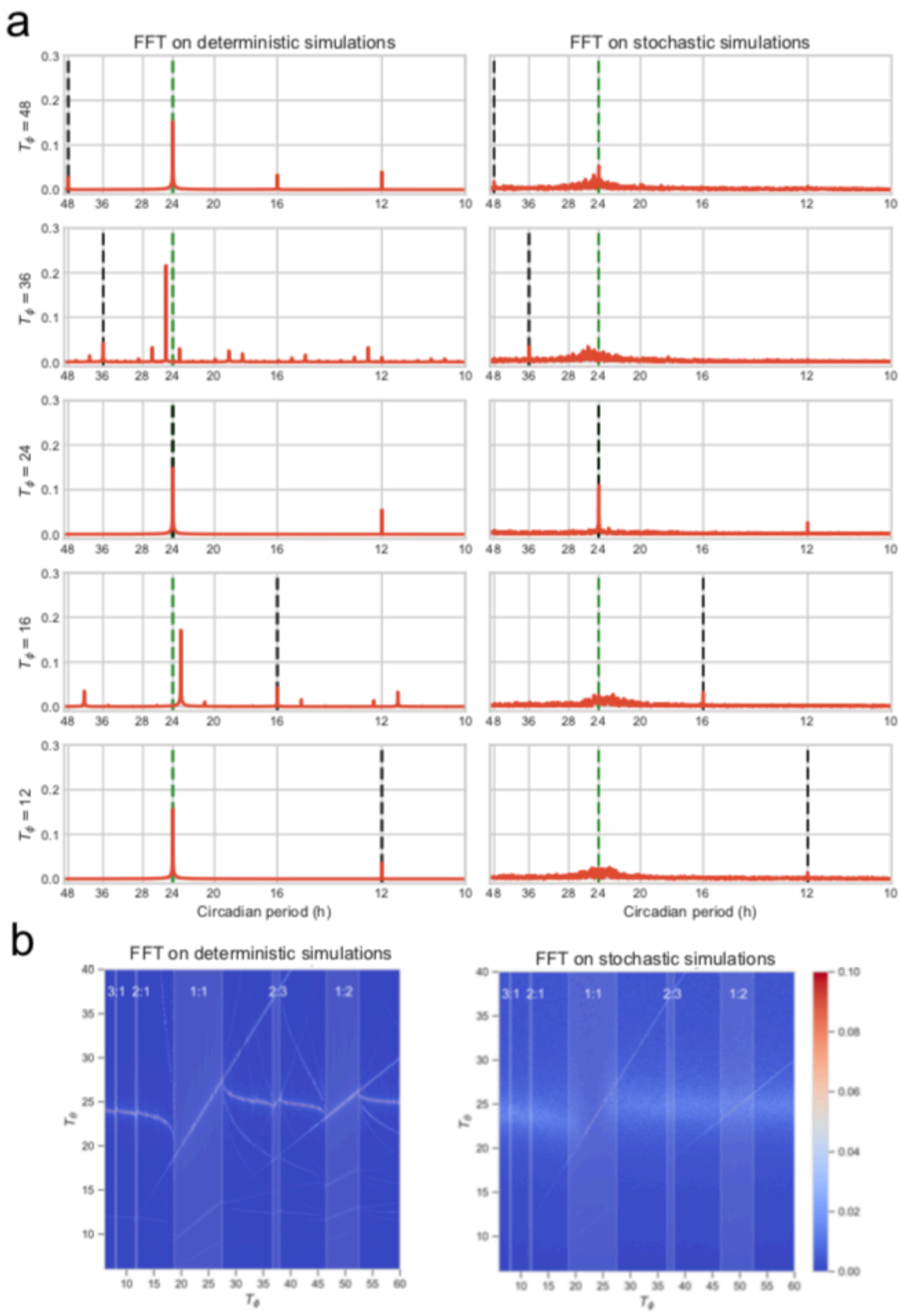


**Supplementary Figure 4: Spectral analysis of simulated traces shows that 1:1 phase-locking is robust against noise.** (**a**) Spectral analysis of long simulated circadian traces ($t_f$=10.000h) using either the deterministic (left, phase diffusion set to zero) or stochastic (right) model, for different cell-cycle periods. Periods of the natural circadian period (24h, green dashed line), and that of the entraining cell cycle (black dashed lines) are indicated. (**b**) Power spectra presented in (a) shown as heatmaps for 350 different cell-cycle periods (see also Movie 3). Phase-locked intervals are observed in the deterministic model (left) as lines for the fundamental and few harmonics. Only 1:1 and the 1:2 are visible in the presence of noise (right).

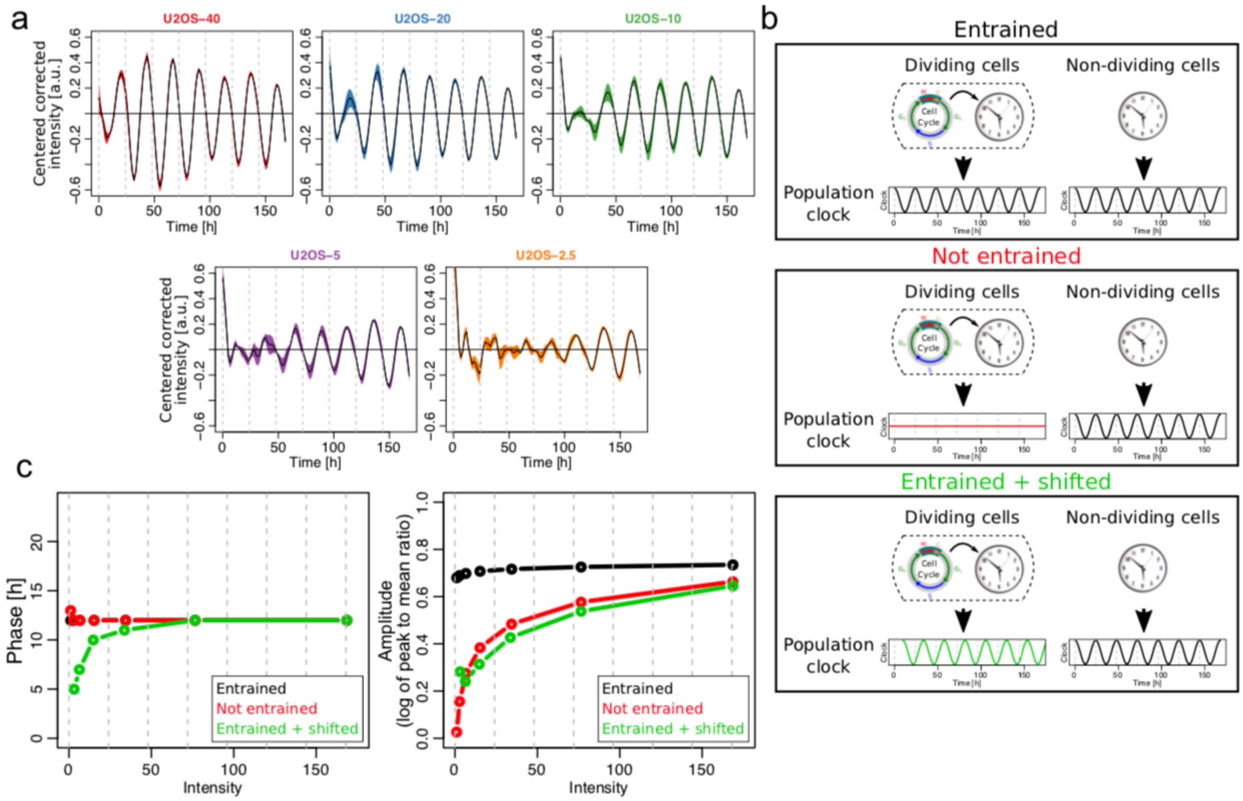


**Supplementary Figure 5: The circadian clock of dividing cells does not entrain to temperature cycles**

(**a**) Averaged (n=6) Bmal1-Luc intensities and 95% confidence intervals from U2OS-Dual Bmal1 luciferase signal centered and corrected for temperature artifact (Supplementary Information). Results were obtained by plating different number of cells (40k, 20k, 10k, 5k, or 2.5k) at the beginning of the experiment. (**b**) Pictograms depicting three different models: i) the circadian oscillators in dividing cells adopt the same circadian profile as non-dividing entrained cells; ii) are not entrained; or iii) are entrained, but with a different phase compared to non-dividing cells. (**c**) Acrophase (left) and amplitude (log of peak to mean ratio) (right) in function of intensity obtained from simulations of the three models in b). Results here should be compared with Figure 6b in the main text.
