## Supplementary information for "Low-dimensional Dynamics of Two Coupled Biological Oscillators"

**4 Analysis of a population of bioluminescence traces under temperature entrainment** **13**

### 1 Reconstruction of the dynamical model

We hypothesized a generic model for the observed fluorescence traces and then optimized its parameters using a maximum likelihood criterion. The main objective is to estimate the coupling function $F(\theta, \phi)$ (expressed in terms of the phases of the two oscillators) representing the influence of the cell-cycle on the circadian clock. The method uses several steps, which are detailed in the following sections.

#### 1.1 Models for the phases and signals

##### 1.1.1 Phase model for dividing cells

The circadian phase is modeled as a diffusion-drift process, while the cell-cycle has simpler, piece-wise linear dynamics between two divisions. This is motivated by previous work where we have shown that the influence of the clock on the cell-cycle was very weak, and probably nonexistent [1]. Therefore we focus here on a precise characterization of the coupling function representing the cell-cycle influence on the clock, and then study the dynamical implications.

We first introduce some notation. $\theta, \phi \in [0; 2\pi[\times[0; 2\pi[$ represent the phases of the circadian clock and the cell-cycle, respectively. The intrinsic period of the circadian clock, $T_\theta$, is kept fixed to $24h$, while the cell-cycle intervals $T_\phi^i$ are indexed on the division-to-division interval $i$. The coupling function $F(\theta, \phi)$ represents the influence of the cell-cycle phase on the circadian clock phase. $\sigma_\theta$ is the noise strength of the circadian phase, the noise itself being modelled through a Wiener process $W_t$. The stochastic phase model is a two-dimensional diffusion drift written as follows:

$$\begin{cases} \mathrm{d}\theta_t = \frac{2\pi}{T_\theta}\mathrm{d}t + F(\theta_t, \phi_t)\mathrm{d}t + \sigma_\theta\,\mathrm{d}W_t \\ \mathrm{d}\phi_t = \frac{2\pi}{T_\phi^i}\mathrm{d}t \end{cases} \quad (1)$$

##### 1.1.2 Phase model for non-dividing cells

In the case of quiescent cells, the model simplifies to:

$$\mathrm{d}\theta_t = \frac{2\pi}{T_\theta}\mathrm{d}t + \sigma_\theta \mathrm{d}W_t \quad (2)$$

##### 1.1.3 Model for the fluorescence signal

The experimental signals obtained from microscopy show noisy oscillations with variations in the amplitude of the maxima as well as in the fluorescence background. For convenience, we centered and rescaled all single cell traces such that the 5th percentile is 0 and 95th percentile is 1.

The phase $\theta_t$ is linked to the signal $S_t$ via a function $w(\theta_t)$, which thus defines the phase in our model. In order to use a common definition of the phase, in particular one that does not depend on temperature or cell type (*i.e.* NIH3T3 and U2OS cells), we estimated a single function $w(\theta_t)$, as the the average of all peak-to-peak signals of non-dividing cells. This showed that indeed NIH3T3 and U2OS cells yield very similar functions, and we therefore used the average as a fixed function for all analyses (Fig. 1).

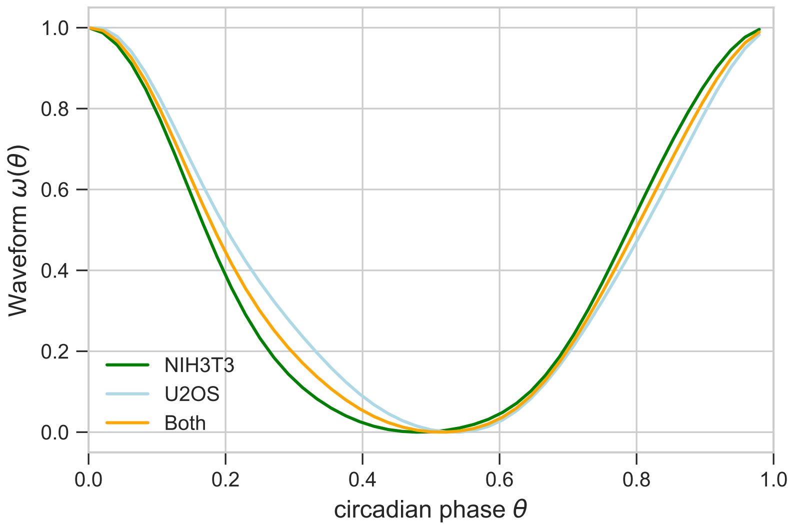

Figure 1: Estimated from NIH3T3 (green) and U2OS (blue) traces reveal little difference between them. To keep a consistent definition for the phase, the final function was taken as the average (yellow).

To take into account the variations of amplitude and background, two Ornstein-Uhlenbeck (O-U) processes are used: $A_t$ and $B_t$.

$$\begin{cases} \mathrm{d}A_t = -\gamma_A(A_t - \mu_A)\mathrm{d}t + \sigma_A\,\mathrm{d}W_t \\ \mathrm{d}B_t = -\gamma_B(B_t - \mu_B)\mathrm{d}t + \sigma_B\,\mathrm{d}W_t \\ S_t = \exp(A_t)w(\theta_t) + B_t + \xi \end{cases} \tag{3}$$

In this parametrization, the stationary mean and variance are $\mathrm{E[X_t]} = \mu_\mathrm{X}$ and $\mathrm{Var[X_t]} = \frac{\sigma^2_\mathrm{X}}{2\gamma_\mathrm{X}}$, for $X = A, B$, respectively. $\xi$ represents additional experimental (measurement) white noise with zero mean and variance $\sigma_e^2$.

This model assumes that the amplitude and the background fluctuations are independent from the phase (see Section 1.3 in this document).

###### 1.1.4 Conversion of the model into a Hidden Markov Model (HMM)

A HMM is defined as a stochastic triplet $\mathbf{\Omega} = \{\boldsymbol{\pi}, \boldsymbol{A}, \boldsymbol{E}\}$, where $\boldsymbol{\pi}$ is the vector containing the initial probability distribution of the modeled Markov process, and the matrices $\mathbf{A}$ and $\mathbf{E}$ contain the transition and emission probabilities of the process [2].

To define the transition and emission matrices, we first discretize the model. The discrete phase, amplitude and background domains are defined as:

$$\begin{cases} \mathbf{\Psi} = \{k\Delta_\Psi | k \in \{0,1,...,N-1\}, \Delta_\Psi = 2\pi/N\} = \{\psi_0, ..., \psi_{N-1}\} \\ \boldsymbol{\mathcal{A}} = \{A_{min} + k\Delta_A | k \in \{0,1,...,M-1\}, \Delta_A = \frac{A_{max}-A_{min}}{M}\} = \{a_0, ..., a_{M-1}\} \\ \boldsymbol{\mathcal{B}} = \{B_{min} + k\Delta_B | k \in \{0,1,...,M-1\}, \Delta_B = \frac{B_{max}-B_{min}}{M}\} = \{b_0, ..., b_{M-1}\} \end{cases}, \tag{4}$$

with $N$ and $M$ the numbers of hidden states for the phase and the O-U processes, respectively.

The maxima ($A_{max}$, $B_{max}$) and minima ($A_{min}$, $B_{min}$) are chosen at least three standard deviations away from the mean of the corresponding O-U processes. The full hidden state space is then $\boldsymbol{\mathcal{X}} = \boldsymbol{\Psi} \times \boldsymbol{\mathcal{A}} \times \boldsymbol{\mathcal{B}}$.

The transition probabilities are then obtained from the following:

$$p(\theta_{t+dt}|\theta_t = \psi_i, \phi_t = \psi_j) = N\left(\psi_i + \frac{2\pi}{T_\theta}dt + F(\psi_i, \psi_j)dt, \sigma_\theta^2 dt\right) \ , \tag{5}$$

where we have made the approximation that $dt$ is small.

For the O-U processes, the results are well-known [3]:

$$p(A_{t+dt}|A_t = a_i) = N\left(\mu_A + (a_i - \mu_A)e^{-\gamma_A dt}, (1 - e^{-2\gamma_A dt})\frac{\sigma_A^2}{2\gamma_A}\right) \ , \tag{6}$$

$$p(B_{t+dt}|B_t = b_i) = N\left(\mu_B + (b_i - \mu_B)e^{-\gamma_B dt}, (1 - e^{-2\gamma_B dt})\frac{\sigma_B^2}{2\gamma_B}\right) \ , \tag{7}$$

All the transitions between the three processes are assumed to be independent:

$$\mathbf{A}_{ijk,lm,no} = p_{tr}(\psi_k|\psi_i, \psi_j)P_{tr}(a_m|a_l)P_{tr}(b_o|b_n) \ . \tag{8}$$

Finally, the probability of the observations obeys to:

$$p(O_t|A_t = a_i, B_t = b_j, \theta_t = \psi_k) = \frac{1}{\sigma_e\sqrt{2\pi}}e^{-\frac{1}{2}\left(\frac{\exp(a_i)w(\psi_k)+b_j-O_t}{\sigma_e}\right)^2} \ . \tag{9}$$

The emission matrix $\boldsymbol{E_t}$ for each time point can then be computed as:

$$\boldsymbol{E}_{t,ijk} = p_e(O_t|a_i, b_j, \psi_k) \ . \tag{10}$$

The fixed cell-cycle phases, given at each time point through a linear interpolation are noted as $\boldsymbol{\Phi}$, such that $\boldsymbol{\Phi} = \{\phi_0, \ldots, \phi_t, \ldots, \phi_T\}$ with $\phi_t \in \boldsymbol{\Psi}\forall t$.

#### 1.2 Parameters of the model

In all analyses, the number of states for the phase, $N$, and for the O-U processes, $M$, were taken as $N = 48$ and $M = 30$, yielding a total number of discrete states of 43200.

##### 1.2.1 Parameters of non-dividing cells

We first discuss the parameters describing the circadian oscillations in individual cells, which are thought to be independent of the coupling with the cell cycle. These parameters concern the oscillator period, phase noise, amplitude and background processes, as well as the experimental noise. The parameters were estimated as described below, and are given in Table 1 for both NIH3T3 and U2OS cells. Since the estimates were found to be very similar at the three experimental temperatures, we considered fixed (temperature-independent) values.

| | $T_\theta(h)$ | $\sigma_\theta(rad.h^{-1/2})$ | $\mu_A$ | $\sigma_A$ | $\mu_B$ | $\sigma_B$ | $\gamma_A(h^{-1})$ | $\gamma_B(h^{-1})$ | $\sigma_e$ |
|---|---|---|---|---|---|---|---|---|---|
| NIH3T3 | 24 | 0.16 | -0.28 | 0.11 | 0.08 | 0.05 | 0.075 | 0.075 | 0.15 |
| U2OS | 24 | 0.18 | -0.45 | 0.15 | 0.06 | 0.05 | 0.075 | 0.075 | 0.15 |

Table 1: Set of parameters for the single cell circadian oscillators in NIH3T3 and U2OS cells. The values of $\sigma_A$, $\sigma_B$ and $\sigma_e$ are in units of the centered and rescaled signals, see section 1.1.3

The circadian oscillator period $T_\theta$ was estimated by averaging the peak to peak times in Rev-Erb$\alpha$-YFP signals on the whole set of non-dividing traces. The resulting value was 24.28h (NIH3T3, whole dataset), rounded for convenience.

To estimate the phase noise $\sigma_\theta$, we used the property that the peak-to-peak time distribution of the circadian phase (modeled as a diffusion-drift process) $\theta$ obeys:

$$T_{2\pi} \sim IG(\mu = \frac{2\pi}{\omega_\theta} = T_\theta, \lambda = \frac{(2\pi)^2}{\sigma_\theta^2}), \tag{11}$$

where $IG(\lambda, \mu)$ stands for the inverse Gaussian distribution with mean $\mu$ and shape parameter $\lambda$. This distribution has variance $\mu^3/\lambda$. Therefore:

$$\sigma_\theta^2 = \frac{Var[T_{2\pi}]4\pi^2}{T_\theta^3}. \tag{12}$$

This expression was used in the NIH3T3 cells. Because we observed only very few non-dividing U2OS cells, we needed to estimate $\sigma_\theta$ from the dividing traces. Since we observed from traces generated *in silico* that the cell-cycle coupling added about 35% of variability in the peak-to-peak distribution, we corrected the value of $\sigma_\theta$ obtained from dividing U2OS cells for this effect.

The means and noise of the O-U processes were estimated from the set of all minima and maxima of the non-dividing traces. More precisely, the mean background was calculated as the average minimum value of the signal, and the mean log amplitude as the average log difference between the maxima and surrounding minima. Similarly, the noise strengths were obtained from the variances of those quantities, using the relationship for the stationary variances: $\sigma_X^2 = 2\gamma_X^2 Var[X]$, for $X = A, B$.

We assumed that the time constants of the $A$ and $B$ processes were slower than the phase fluctuations occurring within one oscillatory cycle, and therefore chose $\gamma_A = \gamma_B = 1/14h^{-1}$. We verified that values of $\gamma_A$ and $\gamma_B$ in the range of $1/5h^{-1}$ to $1/30h^{-1}$ did not lead to major differences in the resulting coupling function.

The noise parameter $\sigma_e$ was set to 0.15. Since the signals were quantile normalized (see section 1.1.3), this corresponds to a relative error of about 15%.

##### 1.2.2 Parameters of the coupling

###### 1.2.2.1 The EM algorithm

Here we introduce the expectation-maximization (E-M) algorithm, which will be used to estimate the coupling function.

Denote the sequence of observations by $\mathbf{O}$, the state space by $\boldsymbol{\mathcal{X}}$, a given sequence of states by $\mathbf{X}$ and the current and updated set of parameters by $\mathbf{\Lambda}'$ and $\mathbf{\Lambda}$, respectively. The $Q$ function of the EM [4] is written:

$$Q(\mathbf{\Lambda}, \mathbf{\Lambda}') = \sum_{\mathbf{X} \in \boldsymbol{\mathcal{X}}} \log(p(\mathbf{O}, \boldsymbol{X}|\mathbf{\Lambda}))p(\boldsymbol{X}|\mathbf{O}, \mathbf{\Lambda}') \ . \tag{13}$$

Here, the sequence $\mathbf{X}$ is composed of states $\mathbf{x}$ such that $\mathbf{X} = \{\mathbf{x}_1, ..., \mathbf{x}_t, ..., \mathbf{x}_T\}$, and these states are themselves composed of three substates for the phase, the amplitude and the background: $\mathbf{x} = (\psi, a, b)$. In our problem, if we define $i_0$ as the index associated with the first state of the sequence $\boldsymbol{X}$, the probability of the observations and the states can be written as a product:

$$p(\mathbf{O}, \boldsymbol{X}|\mathbf{\Lambda}) = \boldsymbol{\pi}_{i_0} \prod_{t=1}^{T} p(\boldsymbol{X}_{t+1}|\boldsymbol{X}_t, \mathbf{\Lambda})p(O_{t+1}|\boldsymbol{X}_{t+1}, \mathbf{\Lambda}) \ . \tag{14}$$

This equation enables to rewrite the function $Q$ as three separated sums:

$$\begin{aligned} Q(\mathbf{\Lambda}, \mathbf{\Lambda}') = &\sum_{\mathbf{X} \in \boldsymbol{\mathcal{X}}} \log(\boldsymbol{\pi}_{i_0})p(\boldsymbol{X}|\mathbf{O}, \mathbf{\Lambda}') + \sum_{\mathbf{X} \in \boldsymbol{\mathcal{X}}} \left( \sum_{t=1}^{T} \log(p(\boldsymbol{X}_{t+1}|\boldsymbol{X}_t, \mathbf{\Lambda})) \right) p(\boldsymbol{X}|\mathbf{O}, \mathbf{\Lambda}') \\ &+ \sum_{\mathbf{X} \in \boldsymbol{\mathcal{X}}} \left( \sum_{t=1}^{T} \log(p(O_{t+1}|\boldsymbol{X}_{t+1}, \mathbf{\Lambda}) \right) p(\boldsymbol{X}|\mathbf{O}, \mathbf{\Lambda}') \ . \end{aligned} \tag{15}$$

This expression readily extends to several traces by adding another sum over the trace indices. Each term can now be optimized individually, enabling to find the optimal set of parameters for the initial condition, the state transitions (which contain the coupling function), and the emissions (cf. 1.1.4).

###### 1.2.2.2 Estimation of the initial condition

Taking the derivative of the first term in Eq (15)with respect to the components of $\boldsymbol{\pi}$ leads to the optimal initial condition:

$$\boldsymbol{\pi}_i = p(\boldsymbol{X}_0 = \mathbf{x}_i|\mathbf{O}, \mathbf{\Lambda}') \tag{16}$$

###### 1.2.2.3 Estimation of the coupling function

The coupling function is parameterized on a grid of $N^2$ parameters, such that $F_{ij}$ corresponds to the coupling for the pair of phases $(\theta_i, \phi_j) \in \mathbf{\Psi}^2$. Due to this high number of parameters, regularization constraints were added. Specifically, the squared norm of the gradient, $||\nabla F_{ij}||^2 = (\frac{F_{i+1,j} - F_{i,j}}{\Delta\psi})^2 + (\frac{F_{i,j+1} - F_{i,j}}{\Delta\psi})^2$ is used to control for smoothness. In addition, we controlled the sparseness of the coupling function using the squared norm. The penalized version of $Q$ is therefore:

$$Q_p(\mathbf{\Lambda}, \mathbf{\Lambda}') = Q(\mathbf{\Lambda}, \mathbf{\Lambda}') - \lambda_1 \sum_{i,j} ||\nabla F_{ij}||^2 - \lambda_2 \sum_{i,j} {F_{ij}}^2 \tag{17}$$

Starting again from Eq. (15) augmented with these new penalization terms yields:

$$\frac{\partial Q_p(\mathbf{\Lambda},\mathbf{\Lambda}')}{\partial F_{kl}} = \overbrace{\frac{\partial}{\partial F_{kl}}\left[\sum_{\mathbf{X}\in\boldsymbol{\mathcal{X}}}\left(\sum_t \log(p(\boldsymbol{X}_{t+1}|\boldsymbol{X}_t,\mathbf{\Lambda}))\right)p(\boldsymbol{X}|\mathbf{O},\mathbf{\Lambda}')\right]}^{E_1} \underbrace{-\frac{\partial}{\partial F_{kl}}\left[\lambda_1\sum_{i,j}||\nabla F_{ij}||^2+\lambda_2\sum_{i,j}F_{ij}^2\right]}_{E_2} \tag{18}$$

The first part of this equation, $E_1$, corresponds to the state transitions, while the second part, $E_2$, corresponds to the penalization. Equating this to zero to find the maxima conditions, and explicitly taking the sequence of cell-cycle states into account, we obtain:

$$\frac{\partial}{\partial F_{kl}}\Bigg[\sum_{i_1,i_2}\sum_{j_1,j_2}\sum_{k_1,k_2}\sum_t \log(p(\theta_{i_2},a_{j_2},b_{k_2}|\theta_{i_1},\phi_t,a_{j_1},b_{k_1},\mathbf{\Lambda})) \\ p(\mathbf{x}_t=(\theta_{i_1},a_{j_1},b_{k_1}),\mathbf{x}_{t+1}=(\theta_{i_2},a_{j_2},b_{k_2})|\mathbf{O},\mathbf{\Lambda}')\Bigg] = E_2 \tag{19}$$

Note here that $\theta_{i_1},\theta_{i_2},a_{j_1},a_{j_2},b_{k_1},b_{k_2}$ are hidden states, for which we infer a distribution of probability with the HMM, while $\phi_t$ is given as an external parameter. Now, we have (cf. eq. 8):

$$\log\left(p(\theta_{i_2},a_{j_2},b_{k_2}|\theta_{i_1},\phi_t,a_{j_1},b_{k_1},\mathbf{\Lambda})\right)) = \log\left(p(\theta_{i_2}|\theta_{i_1},\phi_t,\mathbf{\Lambda})\right) \\ +\log\left(p(a_{j_2}|a_{j_1},\mathbf{\Lambda})\right)+\log\left(p(b_{k_2}|b_{k_1},\mathbf{\Lambda})\right) \ . \tag{20}$$

Since the transitions probabilities for the amplitude and the background do not depend on the coupling function, the remaining sum leads to the marginal joint distribution of phases at time $t$ and $t+1$, and we find:

$$\frac{\partial}{\partial F_{kl}}\left[\sum_{i_1,i_2}\sum_t \log\left(p(\theta_{i_2}|\theta_{i_1},\phi_t,\mathbf{\Lambda})\right)p(\mathbf{x}_t=(\theta_{i_1},.,.),\mathbf{x}_{t+1}=(\theta_{i_2},.,.)|\mathbf{O},\mathbf{\Lambda}')\right] = E_2 \tag{21}$$

Now, $p(\mathbf{x}_t=(\theta_{i_1},.,.),\mathbf{x}_{t+1}=(\theta_{i_2},.,.)|\mathbf{O},\mathbf{\Lambda}')$ doesn't depend on the new coupling function parameters $F_{kl}$, so it can be treated as a multiplicative constant. Defining $\omega_\theta = 2\pi/T_\theta$, this yields:

$$\frac{\partial}{\partial F_{kl}}\Bigg[\sum_{i_1,i_2}\sum_t \log\left(\frac{1}{\sigma_\theta\sqrt{2\pi dt}}e^{-\frac{1}{2}\left(\frac{\theta_{i_2}-(\theta_{i_1}+\omega_\theta dt+F(\theta_{i_1},\phi_t)dt)}{\sigma_\theta^2 dt}\right)^2}\right) \\ p(\mathbf{x}_t=(\theta_{i_1},.,.),\mathbf{x}_{t+1}=(\theta_{i_2},.,.)|\mathbf{O},\mathbf{\Lambda}')\Bigg] = E_2 \tag{22}$$

All the terms that do not depend on $F_{kl} = F(\theta_k,\phi_l)$ are removed by the derivative, which simplifies to:

$$\sum_{i_2} \sum_{\{t|\phi_t=\phi_l\}} \frac{\theta_{i_2} - (\theta_k + \omega_\theta dt + F_{kl}dt)}{\sigma_\theta^2} p(\mathbf{x}_t = (\theta_k, ., .), \mathbf{x}_{t+1} = (\theta_{i_2}, ., .)|\mathbf{O}, \mathbf{\Lambda}') = E_2 \tag{23}$$

$F_{kl}$ can now be isolated, and we can sum over $\theta_{i_2}$ in the denominator:

$$F_{kl} = \frac{-\sigma_\theta^2 E_2 + \sum_{i_2} \sum_{\{t|\phi_t=\phi_l\}} (\theta_{i_2} - (\theta_k + \omega_\theta dt))\, p(\mathbf{x}_t = (\theta_k, ., .), \mathbf{x}_{t+1} = (\theta_{i_2}, ., .)|\mathbf{O}, \mathbf{\Lambda}')}{dt \sum_{\{t|\phi_t=\phi_l\}} p(\mathbf{x}_t = (\theta_k, ., .)|\mathbf{O}, \mathbf{\Lambda}')} \tag{24}$$

From eq. 18, we find:

$$E_2 = \frac{\partial}{\partial F_{kl}} \left[ \lambda_1 \sum_{i,j} (\frac{F_{i+1,j} - F_{i,j}}{\Delta\psi})^2 + (\frac{F_{i,j+1} - F_{i,j}}{\Delta\psi})^2 + \lambda_2 \sum_{i,j} F_{ij}^2 \right] \tag{25}$$

Taking the derivative, and re-injecting into Eq. 24 yields:

$$\begin{aligned} &F_{kl} \left[ dt \sum_{\{t|\phi_t=\phi_l\}} p(\mathbf{x}_t = (\theta_k, ., .)|\mathbf{O}, \mathbf{\Lambda}') + \frac{8\lambda_1\sigma_\theta^2}{\Delta\psi^2} + 2\lambda_2\sigma_\theta^2 \right] \\ &-\frac{2\lambda_1\sigma_\theta^2}{\Delta\psi^2} F_{k-1,l} - \frac{2\lambda_1\sigma_\theta^2}{\Delta\psi^2} F_{k+1,l} - \frac{2\lambda_1\sigma_\theta^2}{\Delta\psi^2} F_{k,l+1} - \frac{2\lambda_1\sigma_\theta^2}{\Delta\psi^2} F_{k,l-1} \\ &= \sum_{i_2} \sum_{\{t|\phi_t=\phi_l\}} (\theta_{i_2} - (\theta_k + \omega_\theta dt))\, p(\mathbf{x}_t = (\theta_k, ., .), \mathbf{x}_{t+1} = (\theta_{i_2}, ., .)|\mathbf{O}, \mathbf{\Lambda}') \end{aligned} \tag{26}$$

For readability, we define the new following quantities:

$$\begin{cases} Q_1 = dt \sum_{\{t|\phi_t=\phi_l\}} p(\mathbf{x}_t = (\theta_k, ., .)|\mathbf{O}, \mathbf{\Lambda}') + \frac{8\lambda_1\sigma_\theta^2}{\Delta\psi^2} + 2\lambda_2\sigma_\theta^2 \\ Q_2 = -\frac{2\lambda_1\sigma_\theta^2}{\Delta\psi^2} \\ Q_{k,l} = \sum_{i_2} \sum_{\{t|\phi_t=\phi_l\}} (\theta_{i_2} - (\theta_k + \omega_\theta dt))\, p(\mathbf{x}_t = (\theta_k, ., .), \mathbf{x}_{t+1} = (\theta_{i_2}, ., .)|\mathbf{O}, \mathbf{\Lambda}') \end{cases} \tag{27}$$

This gives:

$$F_{kl}Q_1 + (F_{k-1,l} + F_{k+1,l} + F_{k,l-1} + F_{k,l+1})Q_2 = Q_{k,l} \tag{28}$$

This is a linear equation for $F_{kl}$. Since Eq. 26 holds $\forall k, l \in \mathbb{N}^2$, this can be rewritten as:

$$\mathbf{A}\mathbf{x} = \mathbf{b} \tag{29}$$

Where $\mathbf{A}$ is a matrix containing the $Q_1$ and $Q_2$ terms, $\mathbf{x}$ is the vector containing the $F_{kl}$ terms and $\mathbf{b}$ the vector containing the $Q_{k,l}$ terms. Due to the regularization, $Q_1$ is always invertible.

###### 1.2.2.4 Regularization parameters

$\lambda_1$ is found using four-fold cross-validation, *i.e.* by splitting the NIH3T3 dataset into four chunks and scanning which $\lambda_1$ value gives the best generalization, *i.e.* maximizes the likelihood of the left-out test traces. The resulting value is $10^{-6}$.

The value of $\lambda_2$ is set according to the following principle. The update expression for the coupling function (when $\lambda_1 = 0$) reads:

$$F_{kl} = \frac{\sum_{i_2} \sum_{\{t|\phi_t=\phi_l\}} (\theta_{i_2} - (\theta_k + \omega_\theta dt))\, p(\mathbf{x}_t = (\theta_k, ., .), \mathbf{x}_{t+1} = (\theta_{i_2}, ., .)|\mathbf{O}, \mathbf{\Lambda}')}{2\sigma_\theta^2 \lambda_2 + dt \sum_{\{t|\phi_t=\phi_l\}} p(\mathbf{x}_t = (\theta_k, ., .)|\mathbf{O}, \mathbf{\Lambda}')} \tag{30}$$

Thus, $\lambda_2$ buffers the sum $dt \sum_{\{t|\phi_t=\phi_l\}} p(\mathbf{x}_t = (\theta_k, ., .)|\mathbf{O}, \mathbf{\Lambda}')$, especially when the latter is small, *i.e.* for the phase-space points which are rarely visited by the cells. Defining $T$ as the total number of time measurements (from all cells), we set:

$$\lambda_2 = \frac{T\lambda_2'}{2\sigma_\theta^2} dt\ . \tag{31}$$

The interpretation is as follow: given a phase-space state that is visited once in T time points, if $\lambda_2' = \frac{1}{T}$ then the corresponding coupling parameter is halved. More visited states lead to more robust coupling parameters, and conversely for less visited states.

In practice, we want to be able to interpret $\lambda_2'$ independently of the total number of time points, and we therefore compute it in units of cell-cycle periods, such that the coupling parameter of a state visited once every 200 cell-cycles is halved, that is:

$$\lambda_2' = \frac{1}{200T_\phi} \tag{32}$$

##### 1.3 Assessment of model assumptions

Our model for the signal $S_t = \exp(A_t)w(\theta_t) + B_t + \xi$ relied on the assumption that the amplitude $A_t$ and background $B_t$ fluctuations were independent of the phase dynamics.

Moreover, we also assumed that phase noise $\sigma_\theta$ was independent of the phase $\theta$, and that the noise $\xi$ was purely additive.

Fig. 2 shows that the expected amplitudes and backgrounds (computed from from $p(\mathbf{O}|\boldsymbol{X})$ using the forward-backward algorithm [2]) are indeed only weakly dependent on the expected phases (top left and top right). In addition, the expected phase diffusion coefficient is independent from the phase (bottom left), and the noise is indeed additive (bottom right).

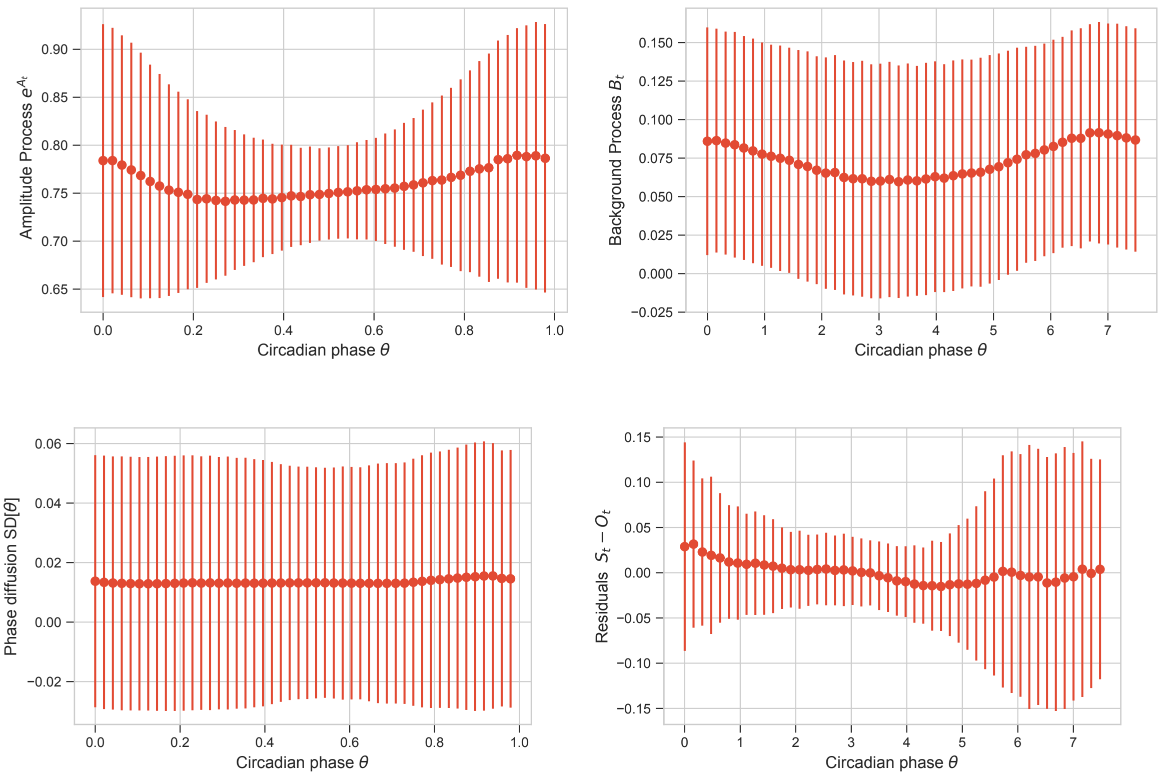

Figure 2: Tests of the assumptions in the model. All the computation are made from the distributions computed on all the traces coming from NIH3T3 cells, all temperatures included. **Top-left:** Mean (over all cells) expected amplitudes (exponentiated) against mean expected phases show that the amplitude is only weakly dependent on the phase. **Top-right:** Mean expected background values against mean expected phases shows that the background is weakly dependent on the phase.**Bottom-left:** Average phase diffusion against mean phase shows independence of the two variables. **Bottom-right:** Model residuals computed from the difference between the expected value of the signal and the experimental data points show that the noise is mainly additive.

#### 1.4 Reliability of the parameter estimation

To assess the reliability of the parameters identified, we generated traces *in silico* and re-estimate the parameters using the same methods as for the experimental traces. The generated traces were of the same scale and length as the experimental traces. The regression parameters $\gamma_A$, $\gamma_B$ and the noise parameters $\sigma_e$ were taken from Table 1.

Table 2 summarizes the results for all estimated parameters, at all temperatures and all cell types. Although we expect some imprecisions due to the stochasticity of the system, the relative error remains low for every parameter.

For the reliability of the estimated coupling function, we refer to the main text (Figure S1, panels b and c).

| | $T_\theta(h)$ | $\sigma_\theta(rad.h^{-1/2})$ | $\mu_A$ | $\sigma_A$ | $\mu_B$ | $\sigma_B$ |
|---|---|---|---|---|---|---|
| Simulated | 24.0 | 0.16 | -0.28 | 0.11 | 0.08 | 0.05 |
| Estimated | 24.0 | 0.16 | -0.24 | 0.11 | 0.04 | 0.05 |

Table 2: Simulated and estimated model parameters.

### 2 Simulations of the dynamical system

#### 2.1 Model

A deterministic model for the phase dynamics is obtained by removing the phase noise term from the full model. In addition, to study the bifurcations (phase locked states) in function of the coupling strength, we added a multiplicative factor for the coupling function called $K$: ($K = 1$ for the biological coupling value).

$$\begin{cases} \dot{\theta} = \frac{2\pi}{T_\theta} + KF(\theta, \phi) \\ \dot{\phi} = \frac{2\pi}{T_\phi} \end{cases} \tag{33}$$

#### 2.2 Phase-locked states

Weakly coupled oscillators can phase-lock when the ratio of their natural period is close to a ratio of integer numbers, *i.e.* $\frac{T_\theta}{T_\phi} \simeq \frac{p}{q}$ with $p, q \in \mathbb{N}$ [5]. To characterize mode-locked states, we estimate $\bar{\omega}_\theta$, defined as the average circadian phase velocity:

$$\bar{\omega}_\theta = \lim_{t\to\infty} \frac{\theta(t)}{t} \ . \tag{34}$$

Phase-locking occurs when $\bar{\omega}_\theta$ remain constant within an interval of cell-cycle frequencies $\omega_\phi$, as represented by Arnold tongue diagrams. Outside of such stable intervals, the dynamics is quasiperiodic.

### 3 Correspondence between cell-cycle phase and biological cell-cycle events

In our model, we assumed a linear progression of the cell-cycle phase between two successive divisions. To get a better handle on the relation between this measure and cell-cycle events, we generated a set of 104 experimental traces from NIH3T3 cells expressing the FUCCI cell-cycle sensor [9]. To obtain estimates of the boundaries for the different cell-cycle events, we normalized and rescaled all fluorescent signals before mapping them to a 0 to $2\pi$ interval (from division to division, Figure 3). Despite biological variability, the growth phase 1 (G1) generally spans from 0 to $0.4 \times 2\pi$ rad, while DNA replication and growth phase 2 (S-G2) usually occur between $0.4 \times 2\pi$ rad and $0.95 \times 2\pi$ rad. Mitosis usually happens from $0.95 \times 2\pi$ rad to $2\pi$ rad.

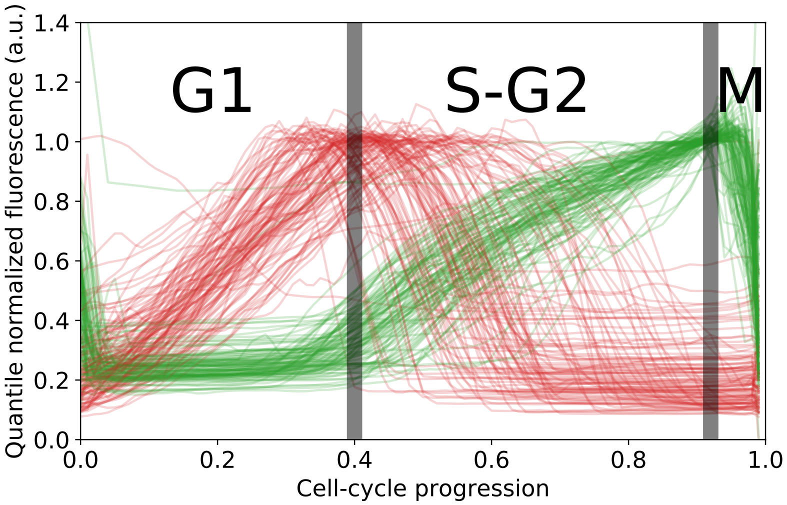

Figure 3: Normalized experimental traces from NIH3T3 cells expressing the FUCCI cell-cycle reporter system enable the association between the physical cell-cycle phase and the biological phase. The red and green fluorescence signals correspond respectively to mKO2-Cdt1 and mAG-Geminin FUCCI reporters. The vertical grey lines denote the (approximate) separation between the different biological cell-cycle phases.

### 4 Analysis of a population of bioluminescence traces under temperature entrainment

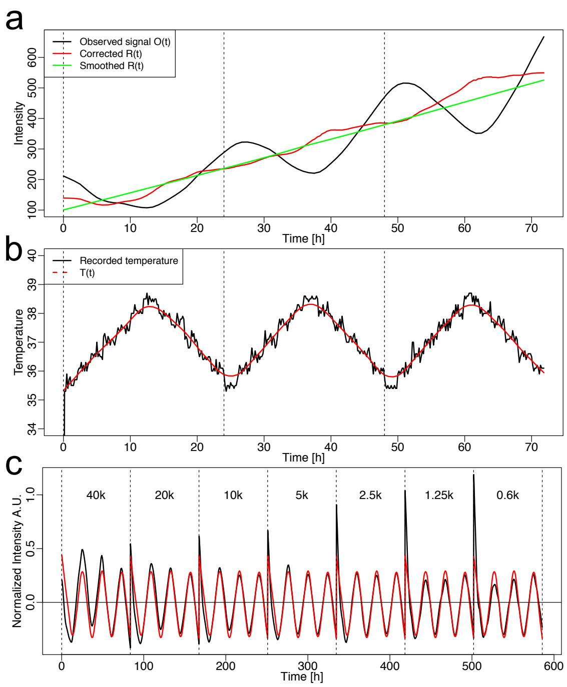

Figure 4: **a:** Observed $O(t)$, smoothed $R(t)$ and corrected $R(t)$ signals obtained from U2OS cells expressing a PGK-luciferase reporter grown at low cell confluence. **b:** Recorded 35.5°C-38.5°C temperature entrainment (black) and smoothed signal (red, $T(t)$) from **a**. **c:** Normalized PGK-Luc signal obtained from U2OS cells grown at different confluences (black) and the optimal fit using $t_d = 80$ min and $k = -0.26$ (red).

The enzymatic activity of luciferase is known to be higher at lower temperature [6]. Since we applied temperature cycles from 35.5°C to 38.5°C for entrainment, even a luciferase reporter driven by a constitutive gene, *e.g. Pgk*, would show an oscillatory signal (Fig. 4, panels a and b)[7, 8]. To correct the signal for this systematic effect, we found that the observed signal $O(t)$ could be well fitted by the following expression

$$O(t) = R(t)(1 + k(T(t - t_d) - T_0) \ , \tag{35}$$

where $R(t)$ is the real signal exempts of any temperature artifact, $T(t)$ is the temperature profile, $T_0 = 37$°C), $k =$ is a magnitude coefficient, and $t_d$ minutes a time delay. To determine the free parameters $t_d$ and $k$, we used the luciferase signal obtained from U2OS cells expressing a PGK luciferase reporter (U2OS-PGK-Luc) which is expected to yield a non-oscillating signal after correction (Fig. 4a). Specifically, we optimized $d$ and $k$ to best fit $O(t)$, after smoothing $O(t)$ to obtain a proxy for $R(t)$. The optimal fit yielded $t_d = 80$ minutes and $k = -0.26$ (Fig. 4c). These values of $t_d$ and $k$ were then used to detrend the circadian luminescence signals using Eq. (35). Importantly, we performed all our luciferase experiments using the Luc2p luciferase (Promega), a destabilized version of the WT Photinus pyralis luciferase optimized for expression in mammals. Consequently, we could use the optimized $t_d$ and $k$ to retrieve the corrected signal $R(t)$ for all our constructs.

### References

[1] Bieler, J., Cannavo, R., Gustafson, K., Gobet, C., Gatfield, D., & Naef, F. (2014). Robust synchronization of coupled circadian and cell cycle oscillators in single mammalian cells. Molecular systems biology, 10(7), 739.

[2] Rabiner, L. R. (1989). A tutorial on hidden Markov models and selected applications in speech recognition. Proceedings of the IEEE, 77(2), 257-286.

[3] Lemons, D. S., & Langevin, P. (2002). An introduction to stochastic processes in physics. JHU Press.

[4] Bilmes, J. A. (1998). A gentle tutorial of the EM algorithm and its application to parameter estimation for Gaussian mixture and hidden Markov models. International Computer Science Institute, 4(510), 126.

[5] Pikovsky, A., Rosenblum, M., Kurths, J., & Kurths, J. (2003). Synchronization: a universal concept in nonlinear sciences (Vol. 12). Cambridge university press.

[6] Koksharov, M.I. and Ugarova, N.N. (2012) Approaches to engineer stability of beetle luciferases. Computational and structural biotechnology journal, 2.

[7] Norrman, K., Fischer, Y., Bonnamy, B., Wolfhagen Sand, F., Ravassard, P. and Semb, H. (2010) Quantitative comparison of constitutive promoters in human ES cells. PloS one, 5.

[8] Qin, J.Y., Zhang, L., Clift, K.L., Hulur, I., Xiang, A.P., Ren, B.Z. and Lahn, B.T. (2010) Systematic comparison of constitutive promoters and the doxycycline-inducible promoter. PloS one, 5.

[9] Sakaue-Sawano, A., Kurokawa, H., Morimura, T., Hanyu, A., Hama, H., Osawa, H., ... & Imamura, T. (2008). Visualizing spatiotemporal dynamics of multicellular cell-cycle progression. Cell, 132(3), 487-498.
